# Multiplex transcriptional characterizations across diverse and hybrid bacterial cell-free expression systems

**DOI:** 10.1101/427559

**Authors:** Sung Sun Yim, Nathan I. Johns, Jimin Park, Antonio L.C. Gomes, Ross M. McBee, Miles Richardson, Carlotta Ronda, Sway P. Chen, David Garenne, Vincent Noireaux, Harris H. Wang

**Affiliations:** Department of Systems Biology, Columbia University, New York, NY, USA; Integrated Program in Cellular, Molecular, and Biomedical Studies, Columbia University, New York, NY, USA; Department of Immunology, Memorial Sloan Kettering Cancer Center, New York, New York, USA; Department of Biological Sciences, Columbia University, New York, NY, USA; School of Physics and Astronomy, University of Minnesota, Minneapolis, MN, USA; Department of Pathology and Cell Biology, Columbia University, New York, NY, USA

**Author notes:** These authors contributed equally (Sung Sun Yim, Nathan I. Johns).

## Abstract

Cell-free expression systems enable rapid prototyping of genetic programs *in vitro*. However, current throughput of cell-free measurements is often limited by the use of single-channel reporter assays. Here, we describe DNA Regulatory element Analysis by cell-Free Transcription and Sequencing (DRAFTS), a rapid and robust *in vitro* approach for multiplexed measurement of transcriptional activities from thousands of regulatory sequences in a single reaction. We employed this method in active cell lysates developed from ten diverse bacterial species. Interspecies analysis of transcriptional profiles from >1,000 diverse regulatory sequences revealed functional differences in gene expression that could be predictively modeled. Finally, we constructed and examined the transcriptional capacities of dual-species “hybrid” cell lysates that can simultaneously harness gene expression properties of multiple organisms. We expect that this cell-free multiplex transcriptional measurement approach will improve genetic circuit prototyping in new bacterial chassis for synthetic biology.

## Introduction

The cell envelope is a key physical barrier that compartmentalizes cellular functions and insulates biological systems from the external environment. However, this barrier also limits direct access to the genome and other cellular components, which can greatly stymie genetic engineering efforts and biomolecular characterizations. Cell-free expression systems have long been established as a simplified *in vitro* approach to overcome these challenges by maintaining cellular processes in the absence of an intact cell membrane^1^. Cell-free systems, made up of either individually reconstituted cellular components^2-4^ or cell lysates^5,6^, can support a variety of catalytic reactions *in vitro* when supplied with energy sources, cofactors and ions^7-9^. These cell-free approaches have facilitated the development of therapeutically useful natural products^10,11^ and biologics with nonstandard amino acids^12^ or chemical moieties otherwise challenging to synthesize^13^. They have also been used to generate viable phage particles directly from purified phage DNA^5,14,15^ and detect pathogens in emerging diagnostics applications^16,17^. Recently, cell-free transcription-translation (TXTL)^18,19^ has increasingly gained popularity as a prototyping platform for synthetic biology^20,21^. By shortcutting time-consuming cloning and transformation steps, TXTL reactions can directly yield proteins and metabolites straight from DNA that encode biosynthetic genes, operons, or pathways. TXTL has also been useful for characterizing synthetic gene circuits with multi-layered components and dynamic behaviors^20,22,23^. However, the correspondence between *in vitro* measurements and actual *in vivo* conditions in live cells remains poorly understood^8,24,25^.

Traditional approaches to track RNA or protein levels in TXTL reactions rely on fluorescent reporters that have limited sensitivity and multiplexing capacity. In contrast, deep sequencing offers a vastly improved approach to characterize thousands of genetic designs or variants simultaneously, which can be produced from pooled DNA synthesis or mutagenesis methods. Such massively parallel reporter assays have enabled detailed studies of governing regulatory features of gene expression, such as 5’ untranslated region (UTR) structure and codon usage, directly in cell populations^26-28^. Cell-free studies have also used similar strategies to study transcription from purified RNA polymerases of phage^29^ and *E. coli*^30^. Large-scale and rapid analysis of genetic components can thus greatly shorten the design-build-test cycle for synthetic biology^31^.

Recently, a number of TXTL systems have been developed from diverse bacterial species^32-35^. These TXTL systems allow better design and testing of genetic circuits and metabolic pathways in new microbes with unique biochemical capabilities^35,36^. Cell-free methods have the potential to more rapidly advance the development of non-model strains that lack characterized genetic parts (i.e. promoters, ribosome binding sites) into useful chassis for industrial synthetic biology. Additionally, the use of diverse TXTLs opens the door for systematic assessment of compatibility between circuit components and various chasses. However, the utility of these *in vitro* approaches is dependent on their correspondence to *in vivo* conditions, which are not well-established.

Here, we describe a new deep sequencing-based multiplex strategy to rapidly measure the activities of thousands of regulatory sequences in a single cell lysate reaction, called <u>D</u>NA <u>R</u>egulatory element <u>A</u>nalysis by cell-<u>F</u>ree <u>T</u>ranscription and <u>S</u>equencing (DRAFTS). We first outline a simple pipeline to rapidly develop and characterize cell-free expression systems in new microbes, which we applied to generate active cell lysates from diverse and industrially useful bacteria. Applying DRAFTS to libraries of natural DNA regulatory sequences, we systematically compared transcriptional measurements made *in vitro* in TXTL versus *in vivo* in cell populations. We then analyzed transcriptional patterns across ten diverse bacteria to identify common and diverging transcriptional capacities between different species as well as in “hybrid” cell lysates made from multiple species. This study provides a guiding roadmap to generate, optimize, and analyze new cell-free expression systems using massively parallel reporter assays to expand the toolbox for genetic prototyping for synthetic biology and biomanufacturing applications.

### Multiplex quantification of 5’ regulatory sequence libraries using DRAFTS

Thanks to recent advances in DNA synthesis, tens to hundreds of thousands of genetic designs can now be cheaply and quickly built in parallel^27,37^. However, these pooled libraries cannot be easily characterized using traditional TXTL reporter assays (e.g. fluorescence), which can often only support single-channel of measurement. As a result, individual genetic circuits need to be separately generated and tested, which adds significant hands-on burden and limits throughput even in 96-well, 384-well or microfluidic formats^35,38^. On the other hand, deep sequencing methods are ideally suited for quantitative measurements of nucleic acid species in pooled reactions. A key constraint of multiplex measurements in pooled reactions, however, is that individual species must not cross-interact with one another. Given these considerations, we designed DRAFTS for quantitative transcriptional analyses of DNA regulatory sequences in pooled TXTL reactions by next-generation sequencing (**Figure 1a**).

**Figure 1.**
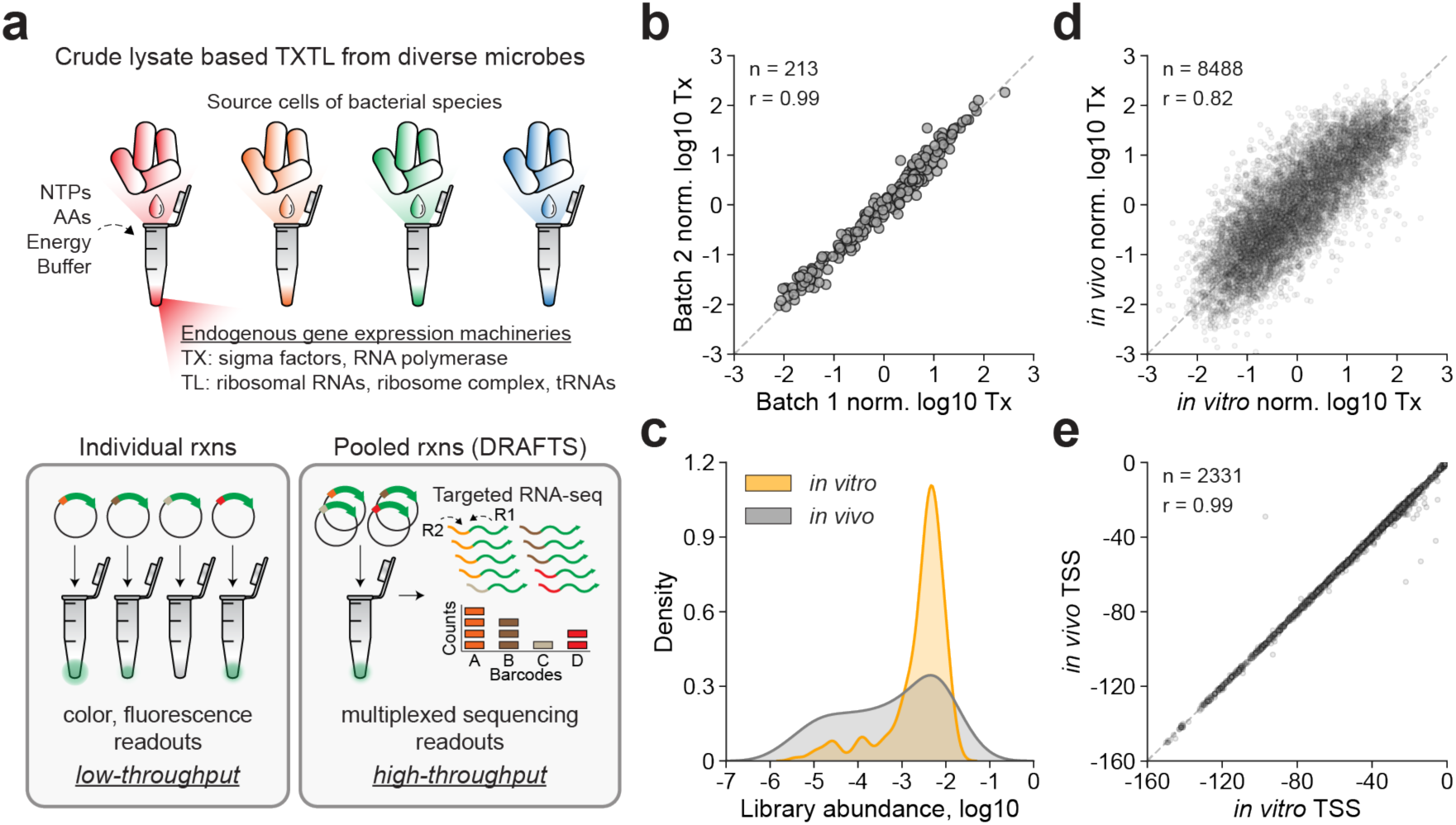
DNA Regulatory element Analysis in Cell-Free Transcription and Sequencing (DRAFTS). **(a)** DRAFTS uses crude cell lysate based cell-free expression systems to harness source host cell’s endogenous gene expression machineries. Compared to conventional single-channel reporters with color or fluorescence readouts, multiplexed sequencing readouts from pooled reactions in DRAFTS scale up the throughput of measurement. **(b)** Biological replicates of transcriptional profiles (Tx) measured in separately prepared cell free extract batches of *E. coli*. **(c)** Comparison of abundances of each library constructs in *in vivo* and *in vitro* measurements. **(d)** Correlation between transcriptional profiles (Tx) for regulatory sequence libraries from *in vitro* and *in vivo* measurements in *E. coli*. **(e)** Correlation between primary TSS calls of regulatory sequences from *in vitro* and *in vivo* measurements in *E. coli*. Sample sizes (n) and Pearson correlation coefficients (r) are shown in each plot. All measurements except **(c)** are based on two technical replicates.

To test whether multiplex transcriptional measurements can be carried out in cell-free expression systems, we first performed pooled *E. coli* TXTL reactions using libraries of DNA regulatory parts (ranging from 234 to 29,250 members) derived from our previous study of natural regulatory sequences^37^ (**Supplementary Data Table 1**). From each TXTL reaction, we performed targeted RNA-seq on specific mRNAs, which were enriched by reverse transcription and 3’-end cDNA ligation of sequencing adaptors (**Suppl. Fig. S1**). The relative RNA abundance of each construct was normalized to its relative DNA abundance, which was determined by DNA-seq of the input regulatory library, to yield the final quantitative measurement of transcription (**see *Methods***). Across different regulatory libraries, transcription levels spanned >5 orders of magnitude and were highly reproducible using the same and separately prepared cell lysates on different days (**Suppl. Fig. S2a, Figure 1b**). Similar to previous reports^37^, transcriptional values were consistent across libraries using alternate N-terminal barcodes and downstream genes (sfGFP or mCherry) (**Suppl. Fig. S2b,c**). We observed that relative transcription levels remained constant across different TXTL reaction times (0.5, 1, 2 or 4 hours) as well as input DNA concentrations (0.025, 0.25, 2.5, or 25 nM) (**Suppl. Fig. S2d,e**).

To compare TXTL transcription levels with *in vivo* values, we transformed each library into the same *E. coli* BL21 strain that was used for cell lysate preparation. We observed that the distribution of DNA libraries was much more uniform *in vitro* than *in vivo*, which is likely due to the decoupling of gene expression from cell fitness *in vitro* and can improve multiplexed measurements (**Figure 1c**). More importantly, both transcription levels and transcription start site positions (TSSs) were highly concordant between *in vitro* and *in vivo* measurements (**Figure 1d,e, Suppl. Fig. S3a-d**). Lastly, the sequences upstream of TSSs were enriched for sigma70 binding sites, indicating an accurate identification of mRNA 5’ positions and enabling compositional analysis of promoter sequences (**Suppl. Fig. S3e,f**). Together, these results demonstrate that *in vitro* TXTL reactions faithfully recapitulate *in vivo* transcription conditions, enabling accurate, more uniform, and multiplex characterization of thousands of regulatory components in a single pooled TXTL reaction.

### A simple pipeline for preparing and optimizing diverse microbial lysates

Given the robustness of multiplex transcriptional measurements in *E. coli* TXTL reactions, we sought to generalize this approach by first generating and optimizing cell-free expression systems from other bacterial species. Recently, several studies have reported TXTL reactions from several non-E. *coli* bacterial species^33-36^. To further expand upon these advances, we established a streamlined experimental pipeline to construct, test and optimize cell lysates from diverse species (***Methods***)^18,19^. In short, cells are grown in rich media, lysed by sonication, incubated in a run-off reaction, and dialyzed (**Figure 2a**)^39^. We used the expression of Broccoli, an RNA fluorescence aptamer, to quantify the transcriptional activity of cell lysates^40^. When bound to the fluorophore DFHBI-1T, Broccoli yields a fluorescence signal that is linearly proportional to its mRNA concentration. We systematically optimized reaction buffer conditions by supplementing with different levels of Mg-glutamate and K-glutamate. The most transcriptionally active buffer conditions were used for all subsequent *in vitro* studies (**Figure 2b**).

**Figure 2.**
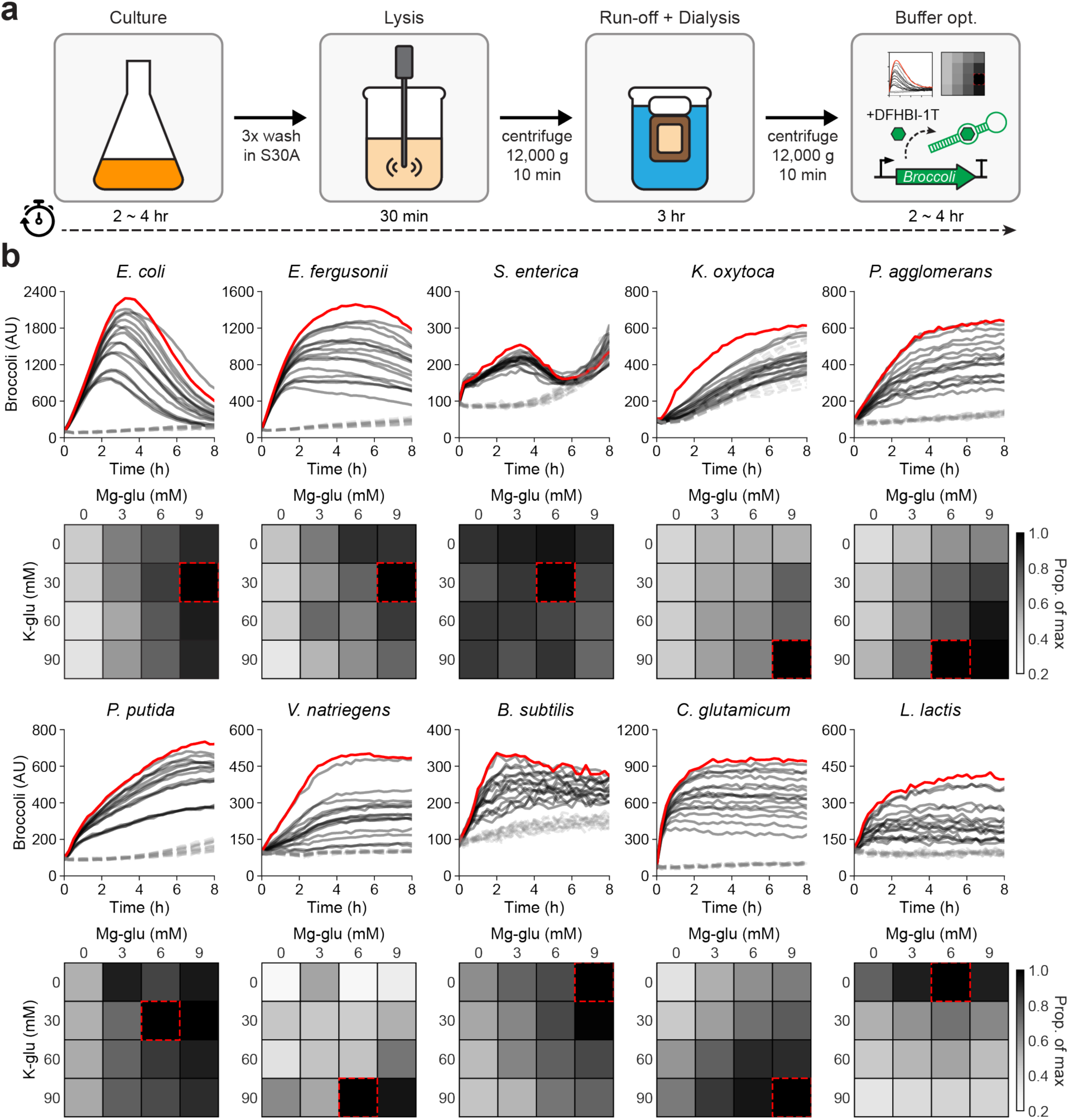
Development of cell-free expression systems of diverse bacterial species. **(a)** Schematic diagram of experimental pipeline for preparation and optimization of cell-free expression systems. **(b)** Optimization of transcriptional output using different concentrations of Mg-glutamate and K-glutamate in 10 bacterial cell-free expression systems with Broccoli as a reporter (solid lines, optimal buffer composition shown in red). No DNA template controls are shown as grey dashed lines. 12.5 nM of DNA template was used in all systems, except *L. lactis* (50 nM used).

In total, we implemented our cell lysate preparation and optimization pipeline on 10 phylogenetically and ecologically diverse bacterial species including seven Proteobacteria (*E. coli, E. fergusonii, S. enterica, P. agglomerans, K. oxytoca, P. putida, V. natriegens*), two Firmicutes (*B. subtilis, L. lactis*) and one Actinobacteria (*C. glutamicum*)*. E. fergusonii* and *P. agglomerans* strains are novel isolates from the murine gut and agricultural waste respectively, while other strains have been previously described. Each strain was first grown in their optimal growth conditions and cell lysates were subsequently prepared using the same general protocol (**Suppl. Methods**). Final cell-free reaction conditions were individually optimized for each species (**Figure 2b** and ***Methods***) and quantified to determine transcriptional yields (**Suppl. Figure 4a**). In the *E. coli* lysate, we found that the expression of the Broccoli reporter peaked after ~4 hours and then decreased thereafter, consistent with previous reports^24^. In contrast, *P. agglomerans, V. natriegens, C. glutamicum* and *L. lactis* showed a prolonged Broccoli signal over time with no observed decrease over 8 hours. These differences could arise as a result of different stability dynamics of the DNA template or resulting mRNA molecules as well as alternative energy recycling processes for sustaining transcription. While transcription requires a few key proteins and cofactors, translation needs many more components with a far greater degree of coordination between them to function properly. Thus, for cell-free lysate reactions, *in vitro* translation is expected to require significantly more tuning and optimization than *in vitro* transcription. Nonetheless, we quantitatively characterized the translational potential of our 10 cell lysates using an eGFP reporter plasmid and detected measurable levels of protein expression in most lysates except for *L. lactis* (**Suppl. Fig S6b**). These results demonstrate that our cell lysate preparation methods are able to generate active cell-free expression reactions from diverse bacteria with relative ease.

**Figure 3.**
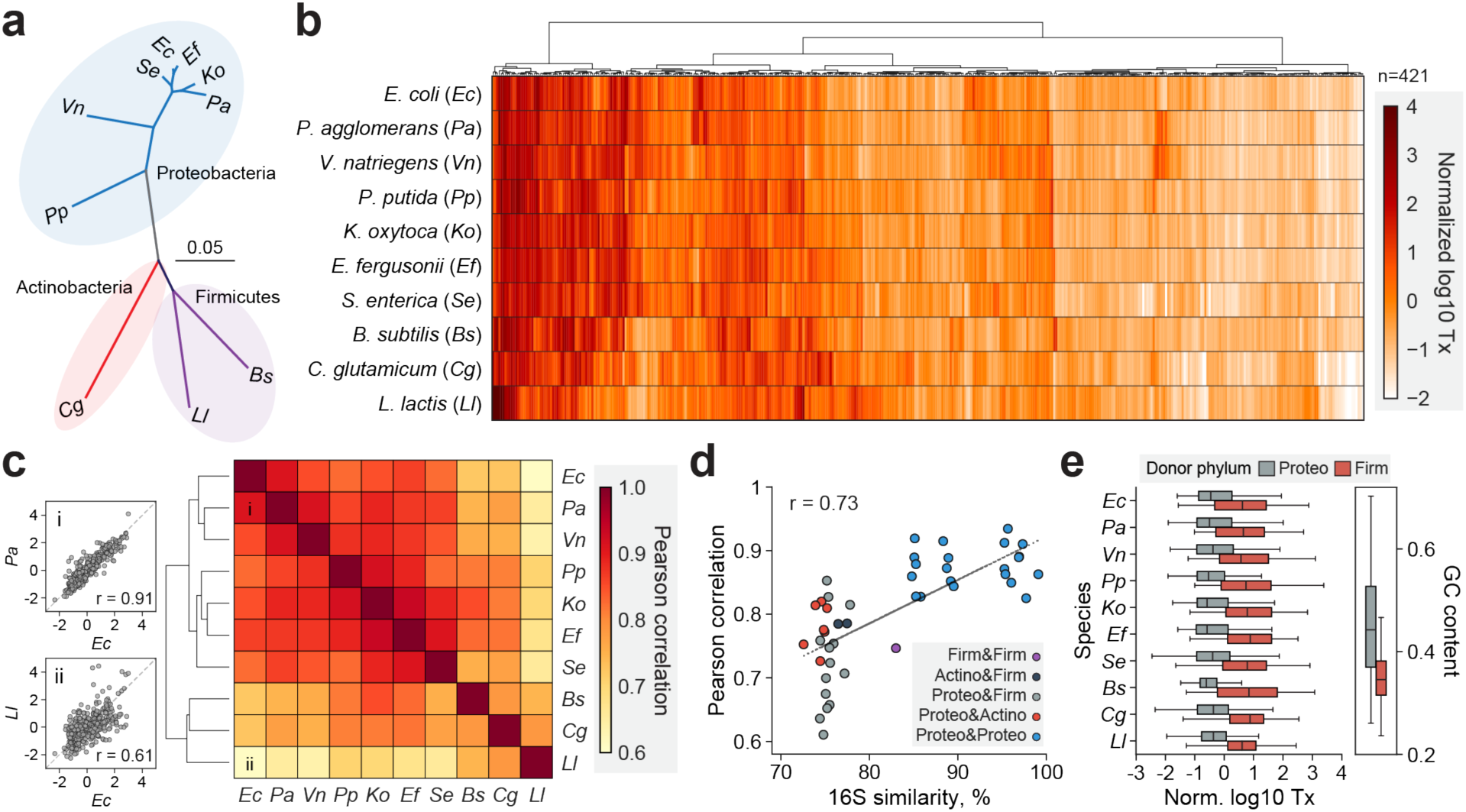
Comparative functional analysis of regulatory sequences across 10 bacterial species through DRAFTS. **(a)** Unrooted phylogenetic tree of 10 bacterial species used in this study based on Jukes-Cantor distance using Neighbor-joining. **(b)** Transcriptional activities (Tx) of 421 regulatory sequences that were active in all 10 bacterial cell-free expression systems. **(c)** Pairwise comparison of transcriptional profiles of 421 universally-active regulatory sequences between bacterial species. Example scatter plots with relatively high and low Pearson correlation in the heatmap (marked i and ii) are shown on left. **(d)** Correlation between evolutionary divergences (16S percent identity) and pairwise Pearson correlation of transcriptional profiles. **(e)** Activity profiles (Tx) of regulatory sequences from donor phyla Proteobacteria and Firmicutes in 10 bacterial species. GC contents of regulatory sequences from the phylogenetic groups are shown on right. All measurements are based on two technical replicates.

**Figure 4.**
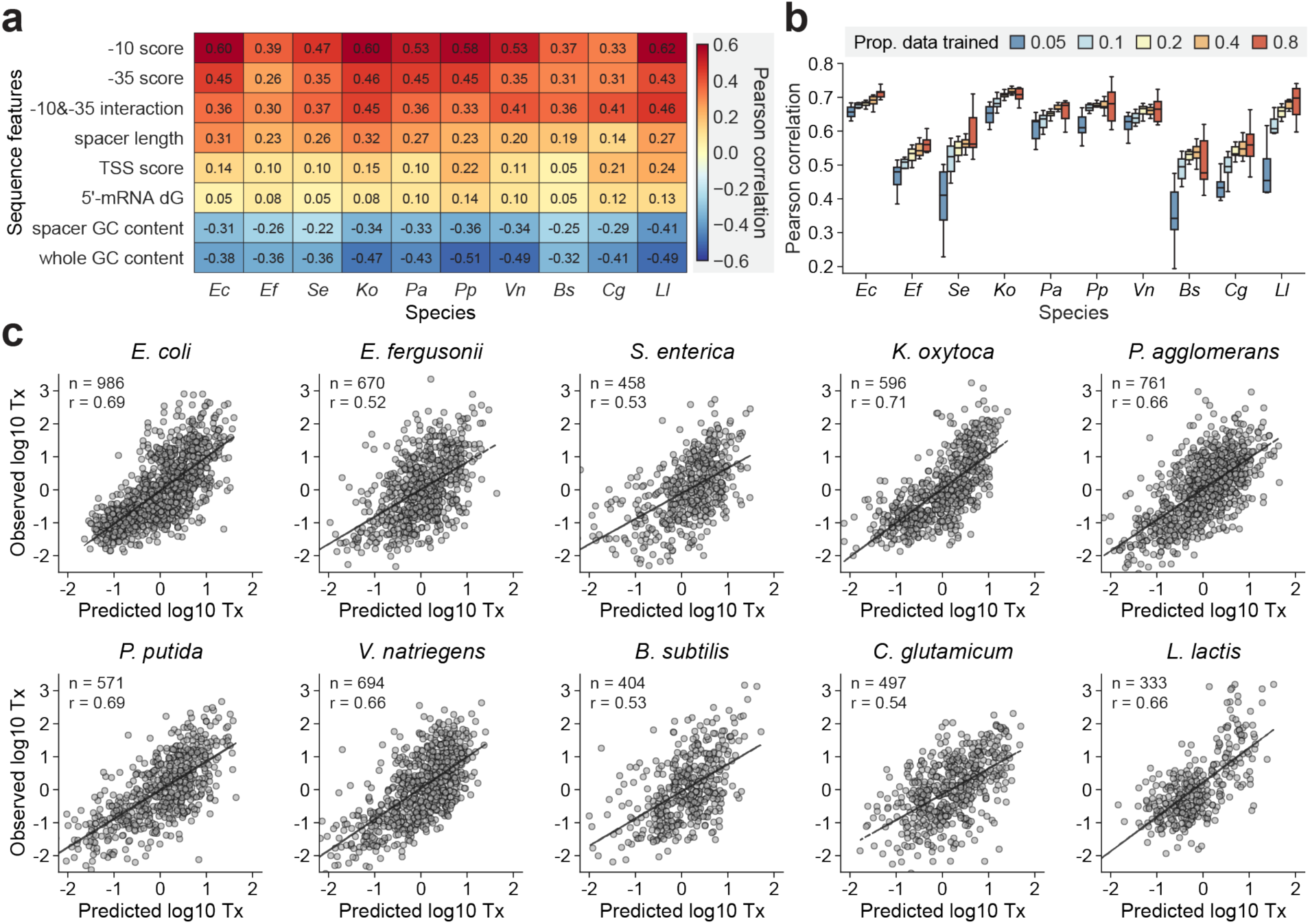
Predictive modeling of transcriptional activation in 10 bacterial species through DRAFTS. **(a)** Correlation between sequence features of regulatory sequences and transcriptional activities in each bacterial species. **(b)** Correlation between predicted and observed transcription levels for each bacterial species with various proportions of data for model training. Data were randomly split for the training and test sets, respectively, and Pearson correlation between predicted and observed transcription levels was computed for 10 times for each proportion. **(c)** Predictive models for transcriptional activation (Tx) in 10 bacterial species using data generated through DRAFTS. Data were randomly split in 10% and 90% for the training and test sets, respectively. All measurements are based on two technical replicates.

To improve the flexibility of our cell-free expression systems, we explored the possible use of uncloned PCR product inputs by attempting to sequester the RecBCD complex, which normally degrades linear DNA templates. Two strategies have been used to enhance the stability of linear DNA involving co-incubation with either GamS from λ-phage or DNA fragments that contain Chi sites^5,41^. To further protect linear DNA templates from RecBCD and other nucleases, we introduced Ku proteins from the non-homologous end-joining (NHEJ) pathway of *Mycobacterium tuberculosis*, which protects damaged DNA by binding to DNA, into our cell-free reactions (**Suppl. Fig. S5a**)^42^. Like GamS and Chi-site DNA, Ku also protected linear DNA templates and enhanced their gene expression in *E. coli* cell-free reactions. Sufficient flanking regions (as little as 100 bp) enabled binding to linear DNA ends by Ku to confer resistance against nuclease degradation (**Supp. Fig. S5b,c**). We then tested Ku in our other cell-free expression systems and found that it improved expression of linear DNA templates in a broad range of species, in contrast to GamS and Chi-site strategies, which worked only in closely related species of *E. coli* and *S. enterica* (**Suppl. Fig. S6**). Together, these results provide a foundation for further optimizations of transcription and translation efficiency in cell-free expression systems and demonstrate their potential for rapid and flexible characterization of genetic designs in new bacterial species.

### Large-scale multiplex transcriptional characterizations in diverse bacteria

Understanding the regulatory capacity of microbes is key for building reliable genetic circuits that experience fewer failures. In previous work, we found that the activity of a regulatory element can vary in different species^37^. A major roadblock in identifying these species-selective regulatory properties is the low efficiency of DNA transformation in many bacteria, which limits multiplex characterization in them. To address this challenge, we first tested a small 234-member regulatory library (RS234)^37^ using DRAFTS in seven species to obtain multiplex transcriptional data. These results were compared with those obtained by transforming the library into each species and measuring *in vivo* transcription levels. Encouragingly, *in vitro* and *in vivo* transcription levels were highly correlated between cell lysates and cell libraries (Pearson’s r between 0.71 and 0.9, **Suppl. Figure S7**), which demonstrates the utility of DRAFTS for accurate multiplex transcriptional measurements in diverse bacteria.

To further scale the throughput of DRAFTS, we mined diverse antibiotic resistance and virulence genes to generate a 1,383-member library of natural regulatory sequences (RS1383) (**Suppl. Fig. S1b** and ***Methods***). Since antibiotic resistance and virulence genes are often mobilized between microbes, we hypothesized that their regulatory regions may exhibit interesting host-range properties. Accordingly, we tested this library in the 10 cell lysates using DRAFTS (**Supplementary Data Table 2**). By using the same RS1383 library in *in vitro* reactions, we can also minimize any confounding contextual effects of different plasmid backbones, which are otherwise needed for *in vivo* characterizations in different species^43^. While the RS1383 library was relatively uniform following *in vitro* reactions in most lysates, a group of sequences (~13%) was significantly depleted in the *L. lactis* lysate. Motif analysis revealed a CCNGG motif in these sequences, which corresponded to a known recognition site of a restriction enzyme (ScrFIR) present in the *L. lactis* genome (**Suppl. Figure S8**)^44^. Therefore, the depletion of these sequences was likely the result of restriction cleavage in *L. lactis* lysates, highlighting the potential interference by bacterial defense systems to confound cell-free studies. We thus removed these sequences from future analyses as they artificially inflated transcription activity calculations.

Many regulatory sequences were transcriptionally active in cell lysates, spanning several orders of magnitude in expression (**Suppl. Fig. 9a**). Hierarchical clustering revealed distinct groups of activity levels across phylogenetically diverse bacteria (**Figure 3a,b**). For each regulatory element, we performed pairwise comparisons of their transcription levels between species (**Figure 3c**, **Suppl. Fig. S9b**). Pearson correlations of these pairwise comparisons showed varying levels of transcriptional concordance, which when clustered further revealed distinct gram-negative and gram-positive groups (**Figure 3c**). Interestingly, more phylogenetically related species shared a more similar transcription profile (Pearson’s r=0.73, **Figure 3d**). Principal component analysis on the RS1383 transcription profiles also revealed two distinct groups for gram-negative and gram-positive species (**Suppl. Fig. S9c**). We did not find distinct patterns of expression across different classes of antibiotic resistance genes across lysates (**Suppl. Figure S10**). However, regulatory sequences derived from Firmicutes, which have AT-rich genomes, tended to be more active overall compared to those from Proteobacteria and other phyla that have higher genomic GC-content (**Figure 3e**), highlighting the importance of regulatory sequence composition. Together, these results provide a higher resolution delineation of the transcriptional capacity and functional similarity between phylogenetically diverse species than that has been described previously.

Finally, we characterized an even larger library of 7,003 regulatory sequences (RS7003)^37^ in the 10 cell lysates (**Supplementary Data Table 3**) to demonstrate the utility of DRAFTS in mechanistic modeling of gene expression. We first examined the relationship between 8 regulatory sequence features known to influence promoter activity (i.e. -10/-35 motif strength and interaction^45^, spacer sequence, TSS region composition, mRNA 5’-end stability, GC%, **Methods**) and the measured *in vitro* expression levels (**Figure 4a**). In line with previous results^37^, the most predictive features for transcriptional activity across species were the strength of the -10 and -35 motifs of sigma70 (positively correlated) and the GC content of the regulatory sequence (negatively correlated). We combined all 8 features to generate a linear regression model of transcriptional activity for each species, using 10% of the dataset for bootstrapped training (**Methods**). These transcriptional models were able to explain up to 50% of the variance in some species (**Figure 4b,c**). Our findings highlight the utility of applying cell-free expression approaches to diverse bacteria for gene regulation characterization and modeling to deepen mechanistic studies^28,45-47^.

### Building synthetic hybrid cell lysates

The diversity of metabolisms and cellular physiologies in the microbial kingdom provides a rich basis to generate cell lysates with unique capacities. Synthetic amalgamation of cell lysates may further extend or broaden capabilities that are otherwise uniquely confined to individual species. With an expanded biochemical repertoire, modified lysates could, in theory, express genes and pathways beyond their native regulatory or metabolic capacities. Furthermore, they could also mimic synthetic bacterial co-cultures, but in an *in vitro* setting that does not require intercellular transport, a limitation in many cross-feeding studies^48^. To test this hypothesis, we synthetically mixed cell lysates from different species into “hybrid lysates” (**Figure 5a**). We first constructed 7 hybrid lysates with equal proportions of two species and examined their ability to produce mRNA and proteins. Interestingly, most hybrid lysates achieved comparable levels of mRNA and protein expression to their single-species counterparts. This result suggests that machineries for gene regulation, transcription, and protein synthesis from evolutionarily distinct species generally do not negatively interfere with one another (**Suppl. Fig. S11, S12**).

**Figure 5.**
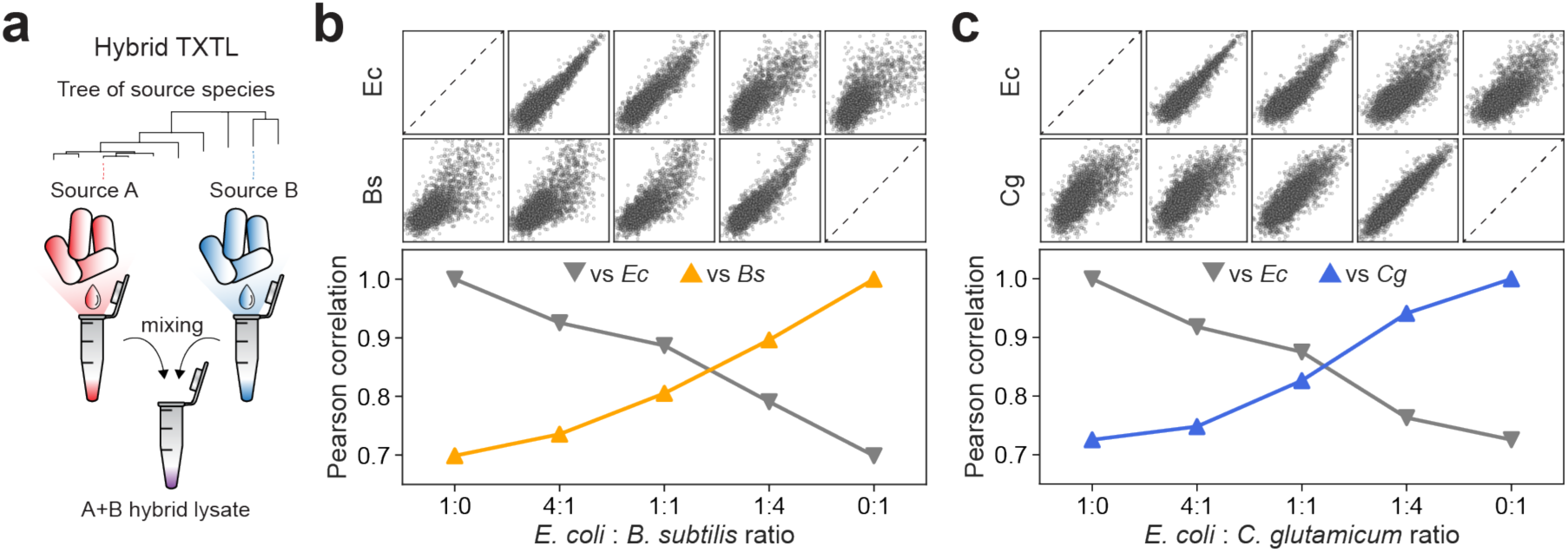
Development and transcriptional characterization of synthetic hybrid TXTL systems. **(a)** Construction of hybrid lysates through extract mixing. Pairwise correlation of RS1383 transcriptional profiles between hybrid lysate and single constituent species lysates for **(b)** *E. coli* + *B. subtilis* and **(c)** *E. coli* + *C. glutamicum* extracts with different mixing ratios. All measurements are based on two technical replicates.

To better delineate the transcriptional properties of hybrid lysates, we then applied DRAFTS using the RS7003 library to examine the transcriptional capacity of *E. coli-B. subtilis* (Ec:Bs) or *E. coli-C. glutamicum* (Ec:Cg) hybrid lysates generated with different mixing ratios (4:1, 1:1, 1:4). Transcriptional profiles from hybrid lysates were compared to one another as well as non-mixed single species (**Figure 5b,c, Supplementary Data Table 4**). Interestingly, equally mixed lysates exhibited transcriptional profiles that were more similar to lysates from both species individually. Tuning the ratio of species in the hybrid lysate resulted in transcriptional profiles more aligned with the majority species (e.g. 1:4 or 4:1) while still retaining some similarity to the minority species. The ability to augment different cell lysates (e.g. Proteobacteria, Firmicutes, Actinobacteria) with properties from functionally different species by simply mixing cell lysates in different ratios and combinations will provide a powerful general approach for *in vitro* synthetic biology. We expect that synthetic hybrid lysates with even higher combinations of species could further broaden the repertoire of biochemical diversity, additively or even multiplicatively.

## Discussion

Cell-free expression systems enable rapid prototyping of natural or synthetic genetic circuits by eliminating time-consuming transformation and cell growth steps. In this study, we developed, optimized, and characterized cell-free lysates for 10 diverse bacterial species from three phyla (Proteobacteria, Firmicutes, Actinobacteria), many of which have never been utilized in cell-free expression reactions. Using DRAFTS in these lysates, we further demonstrated the accurate multiplexed characterization of thousands of DNA regulatory sequences *in vitro*. This deep sequencing-based approach can quantify transcriptional activities across >5 orders of magnitude and simultaneously determine transcriptional start sites. *In vitro* transcriptional measurements faithfully recapitulated *in vivo* expression levels, thus avoiding laborious construction of multiple species-specific libraries and transformation into some challenging or recalcitrant species.

We observed increased regulatory dissimilarity between phylogenetically more distant species, suggesting a functional evolutionary divergence of transcriptional regulation across the natural microbial biome^49^. Differences between microbial species could be exploited to build more refined regulatory circuits that function in specific organisms or multiple targeted species^37^. Using these high-throughput datasets, simple linear models of transcriptional activation could be easily trained to predict promoter strengths in a wide array of organisms. Our identification of a strain-specific restriction enzyme recognition site that affected the stability of DNA templates suggests potential routes for strain optimization in future cell-free studies using diverse bacteria. Advances in cell lysate preparation, especially for organisms requiring anaerobic or fastidious growth conditions, will extend *in vitro* expression systems to new arenas to better understand and improve microbial physiology. Hybrid cell lysates generated by mixing different species could yield *in vitro* systems with new biochemical properties. These synthetic *in vitro* hybrids constitute a step towards harnessing the diverse properties of natural and engineered bacteria into a single *in vitro* reaction, with possible applications in microbiome prototyping, biochemical manufacturing, and the development of artificial minimal cells^48,50^.

While this study focused on characterization of promoters that are constitutively activated, our approach can extend to inducible regulatory systems including riboswitches^51-54^. In these cases, additional regulatory proteins could be co-expressed with a library of regulatory DNA variants in the same *in vitro* reaction and their ligand-response dynamics could be monitored over time. Such approaches could also expedite the characterization of promoters controlled by multiple regulators^47^ and facilitate their integration into complex regulatory networks^51,55^. Transcriptional profiles of entire genetic circuits could also be characterized in TXTL reactions using whole-transcriptome library preparation methods, which have been employed *in vivo* to aid debugging of individual components and dynamic circuit behaviors^56^. Sequencing-based quantification methods for translation such as Ribo-seq^57,58^ could enable *in vitro* multiplex characterization for protein synthesis in different bacteria. Finally, we expect multiplexed sequencing and quantification approaches to also improve analysis of fungal^59^, plant^60^, insect^61^, mammalian^62,63^ and other cell-free expression systems derived from eukaryotic organisms to study and engineer increasingly complex layers of gene regulation using synthetic biology.

## Methods

### Library construction

Regulatory sequences were mined from databases of antibiotic resistance^64^ and virulence genes^65^ (**Suppl. Fig. S1b**). Oligo library design, synthesis, and cloning were performed as previously described^37^. Initial libraries were transformed into *E. coli* MegaX DH10B cells (Invitrogen) and subsequently passaged twice 1:25 in LB+50 μg/mL carbenicillin. Plasmid DNA was extracted (Zymo Midiprep) and column purified once more (PureLink, Invitrogen) to avoid carryover of RNase A into cell-free expression systems. Final concentrations of 10-25 nM were suitable for downstream experiments.

### Lysate preparation

Strain information, media, buffer recipes and growth conditions used for each species can be found in **Supplementary Materials**. *E. fergusonii* and *P. agglomerans* strains were isolated from mouse feces and hemp feedstock (supplied by Ecovative) respectively. Lysates were generated using similar methods used to produce *E. coli* TXTL systems^18,19^. In general, single colonies of each species were inoculated into 4 mL liquid media and grown 6-8 hours. 100-500 μL of the culture was transferred to 50 mL of the same liquid media and grown overnight. 3.3 mL from the overnight culture was transferred to two flasks containing 330 mL growth medium in 1 L baffled flasks (1:100 dilution) and grown to OD_600_ 1.2-1.6. Flasks were rapidly chilled on ice and cells were centrifuged at 4,300xg for 10 minutes at 4°C. Pellets were washed three times with 50 mL S30A buffer in 50 mL centrifuge tubes. The mass of the final pellet was measured and 0.8 mL (1.2 mL for *L. lactis*) S30A buffer was added per gram of pellet mass and then transferred to Eppendorf tubes in 300 uL aliquots. The aliquoted resuspensions were sonicated on ice using a Qsonica125 sonicator with 3.2 mm probe at 35% amplitude for 4 rounds of 30 seconds, with 30-second breaks. Lysates were clarified by centrifuging at 12,000xg for 10 minutes at 4°C. Supernatant was removed and transferred to 2 mL microtubes with screw cap opened. Run-off reactions were performed by incubating clarified lysates at 30°C or 37°C (whichever temperature used for cell growth), 250 rpm for 60-80 minutes. Samples were clarified by centrifuging at 12,000xg for 10 minutes at 4°C. Supernatant was loaded in dialysis cassette (Slide-A-Lyzer 10k MWCO, Thermo Scientific) and dialyzed in S30B buffer for 2-3 hours at 4°C. Samples were clarified once more by centrifuging at 12,000xg for 10 minutes at 4°C, then aliquoted and stored at-80°C.

### Transcriptional Optimization

Transcriptional optimization was performed separately for each species. Cell lysates were combined with amino acids, PEG, and energy buffer as previously described^19^, with additional Mg-glutamate and K-glutamate (both from Sigma Aldrich) at concentrations of 0, 3, 6 or 9 mM and 0, 30, 60, or 90 mM respectively (total 16 combinations), in a skirted white 96-well PCR plate (Bio-rad). A plasmid construct containing strong broad-host range promoter (Gen_18145) identified by our previous study^37^, F30-Broccoli, and B0015 terminator was used as a DNA template, and nuclease-free water was used as a negative control. DNA template (2 uL) and 10 mM DFHBI-1T (0.5 uL, Tocris Bioscience) was added to each well immediately before time course measurements. Fluorescence was tracked for 8 hours using a Synergy H1 plate reader (BioTek) at 30°C using excitation and emission wavelengths of 482 and 505 nm respectively. The concentrations of Mg-glutamate and K-glutamate yielding the highest fluorescence peak was used for future experiments, typically between 1-4 hours.

### Library Transcriptional Measurements

Library measurements were carried out in 10 μL reactions containing 7.5 μL cell-free mixture and 2.5 μL DNA at a final concentration of 10-25 nM. Reactions were incubated at 30°C for 30 min. Each reaction was then split and 2 μL was used for amplification of DNA library and total RNA was extracted from the remaining 5 uL using a Zymo RNA Clean and Concentrator-5 kit, with the remaining volume saved as backups. For *in vivo* measurements, library cultures were grown until OD_600_ ~0.2. RNA was then extracted using RNAsnap^66^. For *in vivo* libraries, cells were lysed using PrepGem bacteria kit (Zygem) for amplification of input DNA library sequences. Unless otherwise stated, RNA sequencing library was prepared by reverse transcription and common adaptor ligation at 3’-end of the cDNA.

Gene-specific reverse transcription was carried out for all RNA samples as follows:

10 μL Total RNA sample (up to 5 μg)
1 μL Reverse transcription primer (20 μM)
1 μL 10 mM dNTPs (Invitrogen)
2.5 μL Nuclease free water

Components were incubated at 65 °C for 5 minutes and chilled on ice for 1 minute. The following were then added to the reaction:

4 μL 5x RT buffer
0.5 μL RNase inhibitor (Ribolock Thermo Scientific)
1 μL Maxima reverse transcriptase (Thermo Scientific)

The reverse transcription reaction was carried out as follows on a 96-well thermocycler (BioRad):

42°C for 90 minutes
50°C for 2 minutes
42°C for 2 minutes
Repeat the two steps above for 9 cycles
85°C for 5 minutes
4°C hold

The completed reaction was incubated with 1 μL RNase H at 37°C for 30 minutes.

Then the common adaptor ligation was carried out as follows:

5 μL cDNA
2 μL Adaptor (40 μM)

Components were incubated at 65 °C for 5 minutes and chilled on ice for 1 minute. The following were then added to the reaction:

2 μL T4 RNA ligase buffer
0.8 μL DMSO
0.2 μL 100 mM ATP
8.5 μL 50% PEG8000
1.5 μL T4 RNA ligase (New England Biolabs)

Reactions were incubated overnight at 22°C for overnight.

Illumina indexes and adaptors were added to both cDNA and input library DNA using a two-step amplification process. All primer sequences are listed in Supplementary Materials. Amplifications were performed using the following PCR reaction mixture and cycling:

10 μL 2x Q5 Hot Start Mastermix (New England Biolabs)
0.2 μL 10x SYBR Green I (Invitrogen)
1 μL Forward primer (10 uM)
1 μL Reverse primer (10 uM)
1 μL 1:10 dilution of library DNA template
6.8 μL Nuclease free water (Ambion)

5 min 98 °C Initial Denaturation
10s 98 °C Denaturation
20s 62 °C Annealing
30s 72 °C Extension
Go to step 2 Until end of exponential amplification
2m 72 °C Final Extension

PCR was performed using a CFX96 Touch Real-Time PCR machine (Bio-Rad). Amplification was stopped as soon as exponential phase ceased, typically around 20 cycles. Samples were diluted 1:100 and amplified again using the same protocol using indexing primers for 8-10 cycles. Samples were then co-purified and examined on a 2% agarose gel to verify correct band sizes of ~ 200 bp for cDNA and 350 bp for input plasmid DNA libraries.

### Read processing and data analysis

After merging paired-end reads using BBmerge^67^ and filtering out mismatched reads, barcode mapping of reads to designed library constructs and expression level calculations were performed by normalizing each construct’s individual RNA abundance by its DNA abundance based on merged counts from two technical replicates, as done in previous studies^27,37^. The log10 transformed expression levels were converted to Z-scores for further comparative analysis across samples.

### Determination of transcription start sites and 5’-end mRNA structure stability

To identify the TSSs of regulatory sequences, alignment of the merged RNA reads with the reference sequence was performed after trimming common adaptor sequence with two random bases at the end. Then we processed TSS calls for each regulatory sequence by an algorithm using kmeans function of scikit-learn package in Python. The algorithm starts with a seed of 16 clusters, then the number of clusters is reduced if two clusters are found within 10 bp of each other or if the cluster contains less than 1% of all reads. Primary TSS was defined as TSS in which >70% of TSS calls lie within the cluster with >200 counts. Other secondary TSSs defined as all the TSSs in which >10% of TSS calls lie. Free energy of 5’-end mRNA structure was computed using the NUPACK package (http://nupack.org)^68^. The 5’-end of mRNA was defined by first 50 bp downstream of TSS of each regulatory sequence. Only promoters with primary TSSs were used for this analysis.

### Motif discovery and scoring

The conserved -10 and -35 motifs of regulatory sequences were identified within the 45 bp preceding primary TSSs using MEME software analysis^69^. Spacer sequences between -35 and -10 regions of promoters were extracted during motif scanning and were given penalty scores for suboptimal sizes deviating up to +/- 2 bp from the optimal 17 bp. We also examined motifs found within the 8 bp of downstream sequences from primary TSS as this region is important for transcription initiation^30,70^. Unless otherwise stated, top 10% highly active promoters were selected for motif analysis. The motif function in the Biopython package was used to scan the aligned sequences by MEME to obtain motif position weight matrix and match scores for each promoter in the regulatory sequence library.

### Statistical methods

All library measurements were performed in duplicate for both RNA and DNA and the Pearson correlation coefficients between most of the biological replicates were >0.9. For most analyses, a cutoff of 15 DNA reads was set to minimize noise from low abundance constructs. All statistics were performed using commonly used Python libraries such as Numpy, Scipy, Pandas, and Seaborn. Linear modeling was performed using the LinearRegression function in scikit-learn package, by randomly splitting the data in 10% and 90% for the training and test sets, respectively, unless otherwise stated.

## Acknowledgements

We thank members of the Wang lab for advice and comments on the manuscript. H.H.W. acknowledges funding support from NSF (MCB-1453219), NIH/NIGMS (U01GM110714-01A1, 1R01AI132403-01), DARPA (HR0011-17-C-0068), DoD ONR (N00014-15-1-2704), the Sloan Foundation (FR-2015-65795). V.N. acknowledges funding support from DoD ONR (N00014-13-1-0074) and Human Frontier Science Program (RGP0032/2015). S.S.Y. thanks support from Basic Science Research Program through the National Research Foundation of Korea funded by the Ministry of Education (NRF-2017R1A6A3A03003401). N.I.J. is supported by an NSF Graduate Research Fellowship (DGE-1644869). We also thank K.J.Jeong (KAIST, Daejeon, Korea) for providing pCES208 plasmid.

## Author contributions

S.S.Y., N.I.J., V.N. and H.H.W. developed the initial concept. A.L.G., C.R., S.P.C., R.M., M.R. and D.G. provided key biological reagents, strains and code. S.S.Y., N.I.J., and J.P. performed experiments and analyzed the results under the supervision of H.H.W.; N.I.J., S.S.Y. and H.H.W. wrote the mansucript with input from all authors.

## Supplementary Materials for Multiplex transcriptional characterizations across diverse and hybrid bacterial cell-free expression systems

**This PDF file includes:**

Methods

Tables S1 to S5

Figs. S1 to S12

## Methods

### Production and purification of Ku in *E. coli*

Ku gene from *Mycobacterium tuberculosis* was synthesized (IDT) and cloned into pET28c with riboJ (BBa_K1679038) and RBS (BBa_B0034) to yield pET28c-Ku. The pET28c-Ku plasmid was introduced to *E. coli* BL21(DE3). Ku was produced in the strain by 0.2 mM IPTG (isopropyl-b-D-thiogalactopyranoside) induction at 18°C for 16 hours. Soluble lysate of the culture was prepared by sonication on ice using a Qsonica125 sonicator with 3.2 mm probe at 40% amplitude for 50 rounds of 30 seconds, with 30 second breaks and clarification by centrifuging at 12,000xg for 10 minutes at 4°C. Recombinant Ku protein in the soluble lysate was purified using Nickel-NTA-agarose. Ku was recovered in the 300 mM imidazole eluates. Eluted Ku protein was dialyzed in PBS for 2-3 hours at 4°C, and stored at -20°C.

Template-switching adaptor ligation

Data in Figure S7 was prepared by Template-switching, an alternative method for addition of common adaptor sequences to cDNA using the following reverse transcription reaction:

5 μL Total RNA sample (up to 5 μg)
1 μL Reverse transcription primer (20 μM)
1 μL 10 mM dNTPs (Invitrogen)
6.5 μL Nuclease free water

4 μL 5x RT buffer
0.5 μL RNase inhibitor (Ribolock Thermo Scientific)
1 μL Maxima reverse transcriptase (Thermo Scientific)
1 μL Template switching oligo (20 μM)

The reverse transcription reaction was carried out as follows on a 96-well thermocycler (BioRad):

The completed reaction was incubated with 1 μL RNase H at 37°C for 30 minutes.

**Table S1.** Bacterial species used in this study and their growth conditions

| **Species** | **Strain (Source)** | **Medium** | **Temp.** | **Aeration** |
| --- | --- | --- | --- | --- |
| *E. coli* | BL21 | 2xYT+P | 37°C | Shaking (220 rpm) |
| *E. fergusonii* | Isolate (Murine feces) | 2xYT+P | 37°C | Shaking (220 rpm) |
| *S. enterica* | Serovar Typhi Ty2 | 2xYT+P | 37°C | Shaking (220 rpm) |
| *K. oxytoca* | M5A1 (DSM 7342) | 2xYT+P | 30°C | Shaking (220 rpm) |
| *P. agglomerans* | Isolate (Agricultural waste) | 2xYT+P | 30°C | Shaking (220 rpm) |
| *P. putida* | KT2440 (ATCC 47054) | 2xYT+P | 30°C | Shaking (220 rpm) |
| *V. natriegens* | ATCC 14048 | BHI+v2 salt | 37°C | Shaking (220 rpm) |
| *B. subtilis* | BD3182 (168 derivative) | 2xYT+P | 30°C | Shaking (220 rpm) |
| *C. glutamicum* | ATCC 13032 | BHI | 30°C | Shaking (220 rpm) |
| *L. lactis* | ATCC 11454 | MRS | 37°C | Static |

**Table S2.** Composition of media, buffer and TXTL components used in this study

| **Medium** | **Composition** |
| --- | --- |
| 2xYT+P | 10 g/L Yeast extract (BD, 288620), 16 g/L Tryptone (BD, 211705), 5 g/L NaCl, 40 mM Potassium phosphate dibasic, 22 mM Potassium phosphate monobasic |
| BHI | 37 g/L BHI (Brain Heart Infusion) (BD, 237500) |
| BHI+v2 salts | 37 g/L BHI (Brain Heart Infusion) (BD, 237500) + v2 salts (204 mM NaCl, 4.2 mM KCl, 23.14 mM MgCl<sub>2</sub>) [Weinstock et al., 2016. Nat. Methods 13:849-51] |
| MRS | 55 g/L MRS (BD, 288130) |

| **Buffer** | **Composition** |
| --- | --- |
| S30A* | 14 mM Mg-glutamate, 60 mM K-glutamate, 50 mM Tris base (pH 7.7 - titrate with acetic acid) + supplement 1 M DTT sol to make 2 mM DTT concentration before use |
| S30B | 14 mM Mg-glutamate, 60 mM K-glutamate, 5 mM Tris base (pH 8.2 - titrate with 2 M Tris base solution) + supplement 1 M DTT sol to make 1 mM DTT concentration before use |

\*[Sun et al., 2013. *J Vis Exp* **79**, e50762]

| **TXTL components** | **Composition** |
| --- | --- |
| Amino acids* | RTS Amino Acid Sampler kit (6 mM of each amino acid, except leucine which is 5 mM) |
| Energy solution (PGA)* | 700 mM HEPES (pH 8), 21 mM ATP, 21 mM GTP, 12.6 mM CTP, 12.6 mM UTP, 2.8 mg/mL tRNA, 3.64 mM CoA, 4.62 mM NAD, 10.5 mM cAMP, 0.95 mM Folinic acid, 14 mM Spermidine + 420 mM 3-PGA |
| Energy solution (PEP) | 700 mM HEPES (pH 8), 21 mM ATP, 21 mM GTP, 12.6 mM CTP, 12.6 mM UTP, 2.8 mg/mL tRNA, 3.64 mM CoA, 4.62 mM NAD, 10.5 mM cAMP, 0.95 mM Folinic acid, 14 mM Spermidine + 420 mM PEP |

\*[Sun et al., 2013. *J Vis Exp* **79**, e50762]

**Table S3.** Composition of cell-free systems for each bacterial species

| **Component** | **Stock conc.** | **Final conc.** | **Volume\*** |
| --- | --- | --- | --- |
| Lysate** | 1 | 0.33** | 28.5 µL |
| Mg-glutamate | 500 mM | 9 | 1.54 µL |
| K-glutamate | 3 M | 30 | 0.86 µL |
| Amino Acids | 6 mM each*** | 1.5 mM each | 21.38 µL |
| Energy Solution (PGA) | 14x | 1x | 6.11 µL |
| PEG-8000 | 40% | 2% | 4.28 µL |
| Water |  |  | 1.47 µL |

*E. fergusonii*
| **Component** | **Stock conc.** | **Final conc.** | **Volume\*** |
| --- | --- | --- | --- |
| Lysate** | 1 | 0.3** | 25.71 µL |
| Mg-glutamate | 500 mM | 9 | 1.54 µL |
| K-glutamate | 3 M | 30 | 0.86 µL |
| Amino Acids | 6 mM each*** | 1.5 mM each | 21.38 µL |
| Energy Solution (PGA) | 14x | 1x | 6.11 µL |
| PEG-8000 | 40% | 2% | 4.28 µL |
| Water |  |  | 4.26 µL |

*S. enterica*
| **Component** | **Stock conc.** | **Final conc.** | **Volume\*** |
| --- | --- | --- | --- |
| Lysate** | 1 | 0.33** | 28.17 µL |
| Mg-glutamate | 500 mM | 6 | 1.03 µL |
| K-glutamate | 3 M | 30 | 0.86 µL |
| Amino Acids | 6 mM each*** | 1.5 mM each | 21.38 µL |
| Energy Solution (PGA) | 14x | 1x | 6.11 µL |
| PEG-8000 | 40% | 2% | 4.28 µL |
| Water |  |  | 2.32 µL |

\*enough for ~8 single reactions
\*\*input as volumetric ratio decided based on total protein conc. of the lysate\*
\*\*\*except Leucine which is 5 mM

**Table S3. (Continued)***K. oxytoca*
| **Component** | **Stock conc.** | **Final conc.** | **Volume\*** |
| --- | --- | --- | --- |
| Lysate** | 1 | 0.3** | 25.79 µL |
| Mg-glutamate | 500 mM | 9 | 1.54 µL |
| K-glutamate | 3 M | 90 | 2.57 µL |
| Amino Acids | 6 mM each*** | 1.5 mM each | 21.38 µL |
| Energy Solution (PGA) | 14x | 1x | 6.11 µL |
| PEG-8000 | 40% | 2% | 4.28 µL |
| Water |  |  | 2.47 µL |

*P. agglomerans*
| **Component** | **Stock conc.** | **Final conc.** | **Volume\*** |
| --- | --- | --- | --- |
| Lysate** | 1 | 0.28** | 23.52 µL |
| Mg-glutamate | 500 mM | 6 | 1.03 µL |
| K-glutamate | 3 M | 90 | 2.57 µL |
| Amino Acids | 6 mM each*** | 1.5 mM each | 21.38 µL |
| Energy Solution (PGA) | 14x | 1x | 6.11 µL |
| PEG-8000 | 40% | 2% | 4.28 µL |
| Water |  |  | 5.26 µL |

**Table S3. (Continued)***V. natriegens*
| **Component** | **Stock conc.** | **Final conc.** | **Volume\*** |
| --- | --- | --- | --- |
| Lysate** | 1 | 0.31** | 26.23 µL |
| Mg-glutamate | 500 mM | 6 | 1.03 µL |
| K-glutamate | 3 M | 90 | 2.57 µL |
| Amino Acids | 6 mM each*** | 1.5 mM each | 21.38 µL |
| Energy Solution (PGA) | 14x | 1x | 6.11 µL |
| PEG-8000 | 40% | 2% | 4.28 µL |
| Water |  |  | 2.55 µL |

*B. subtilis*
| **Component** | **Stock conc.** | **Final conc.** | **Volume\*** |
| --- | --- | --- | --- |
| Lysate** | 1 | 0.32** | 27.67 µL |
| Mg-glutamate | 500 mM | 9 | 1.54 µL |
| K-glutamate | 3 M | 0 | 0 µL |
| Amino Acids | 6 mM each*** | 1.5 mM each | 21.38 µL |
| Energy Solution (PGA) | 14x | 1x | 6.11 µL |
| PEG-8000 | 40% | 2% | 4.28 µL |
| Water |  |  | 3.16 µL |

*C. glutamicum*
| **Component** | **Stock conc.** | **Final conc.** | **Volume\*** |
| --- | --- | --- | --- |
| Lysate** | 1 | 0.33** | 28.5 µL |
| Mg-glutamate | 500 mM | 9 | 1.54 µL |
| K-glutamate | 3 M | 90 | 2.57 µL |
| Amino Acids | 6 mM each*** | 1.5 mM each | 21.38 µL |
| Energy Solution (PGA) | 14x | 1x | 6.11 µL |
| PEG-8000 | 40% | 2% | 4.28 µL |
| Water |  |  | 0 µL |

\*enough for ~8 single reactions
\*\*input as volumetric ratio decided based on total protein conc. of the lysate
\*\*\*except Leucine which is 5 mM

**Table S3. (Continued)***L. lactis*
| **Component** | **Stock conc.** | **Final conc.** | **Volume\*** |
| --- | --- | --- | --- |
| Lysate** | 1 | 0.33** | 28.5 µL |
| Mg-glutamate | 500 mM | 6 | 1.03 µL |
| K-glutamate | 3 M | 0 | 0 µL |
| Amino Acids | 6 mM each*** | 1.5 mM each | 21.38 µL |
| Energy Solution (PEP) | 14x | 1x | 6.11 µL |
| PEG-8000 | 40% | 2% | 4.28 µL |
| Water |  |  | 2.84 µL |

\*enough for ~8 single reactions
\*\*input as volumetric ratio decided based on total protein conc. of the lysate
\*\*\*except Leucine which is 5 mM

**Table S4.** Plasmids and libraries used in this study

| Plasmid | Origin | Antibiotic | Description |
| --- | --- | --- | --- |
| TOPO-F30-Broccoli | pBR322 | Carbenicillin | P<sub>Gen_18145</sub>-F30-Broccoli* |
| p70a-deGFP | pBR322 | Carbenicillin | P<sub>70a</sub>-deGFP** |
| pET28c-Ku | pBR322 | Kanamycin | P<sub>T7</sub>-Ku from *M. tuberculosis* |

\*[Filonov et al., 2014. *J Am Chem Soc* **136**, 16299-308]
\*\*[Sun et al., 2013. *J Vis Exp* **79**, e50762]

| Library | Backbone | Origin | Antibiotic | Related Figures |
| --- | --- | --- | --- | --- |
| RS234 | pNJ1 | p15A | Carbenicillin | Fig. 1B, 1C,<br>Fig. S2A, C, D, E, Fig. S7 |
| RS29250 | pNJ1 | p15A | Carbenicillin | Fig. 1D, 1E,<br>Fig. S2B, Fig. S3 |
| RS1383 | pNJ1 | p15A | Carbenicillin | Fig. 3, Fig. S1,<br>Fig. S8, Fig. S9, Fig. S10, |
| RS7003 | pNJ1 | p15A | Carbenicillin | Fig. 4, Fig. 5 |
| RS234 for Ko, Vn | pNJ7 | p15A | Chloramphenicol | Fig. S7 |
| RS234 for Pp | pNJ3 | pBBR1 | Carbenicillin | Fig. S7 |
| RS234 for Bs | pNJ2 | integrative,<br>pBR322 | Chloramphenicol | Fig. S7 |
| RS234 for Cg | pNJ8 | pCG1,<br>p15A | Kanamycin | Fig. S7 |

**Table S5.**
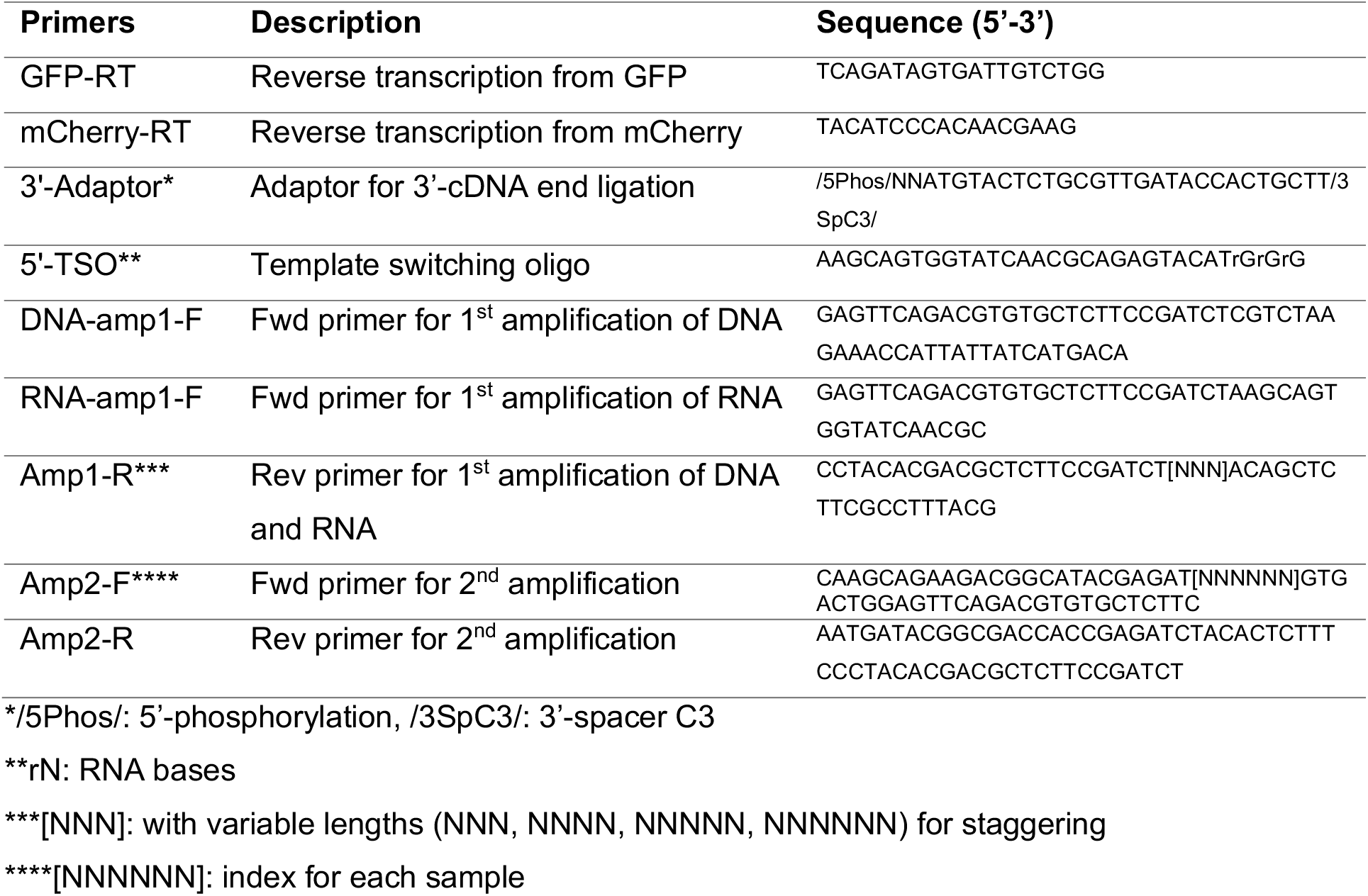
Primers used in this study

**Supplementary Figure S1.**
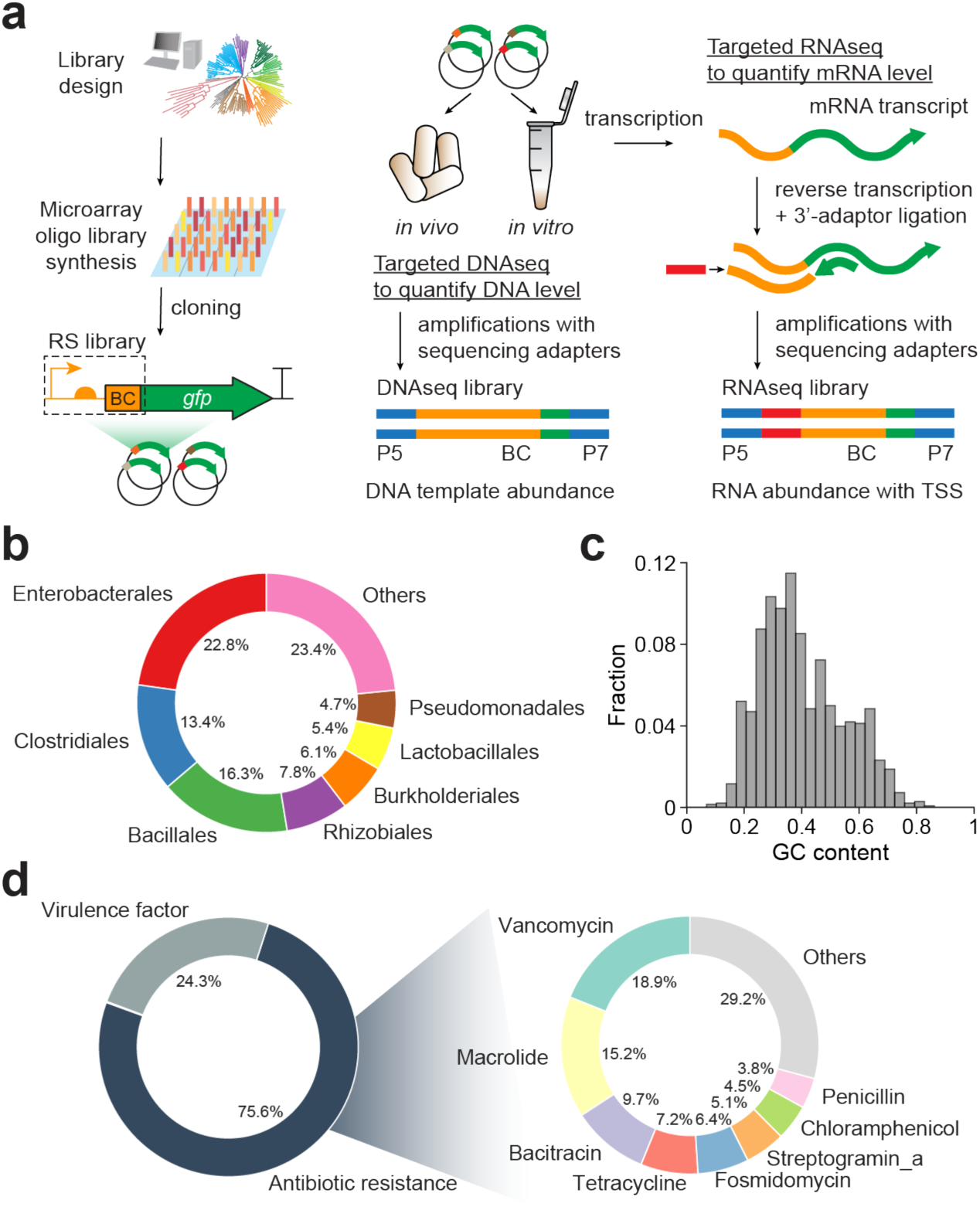
DNA regulatory sequence analysis in cell-free transcription systems. **(a)** Microarray synthesis, library cloning, and high-throughput characterization of regulatory sequences using targeted sequencing of RNA and DNA. **(b)** Phylogenetic origins, **(c)** GC content distribution, **(d)** functional categories of genes where the sequences mined from and related antibiotics of RS1383 library.

**Supplementary Figure S2.**
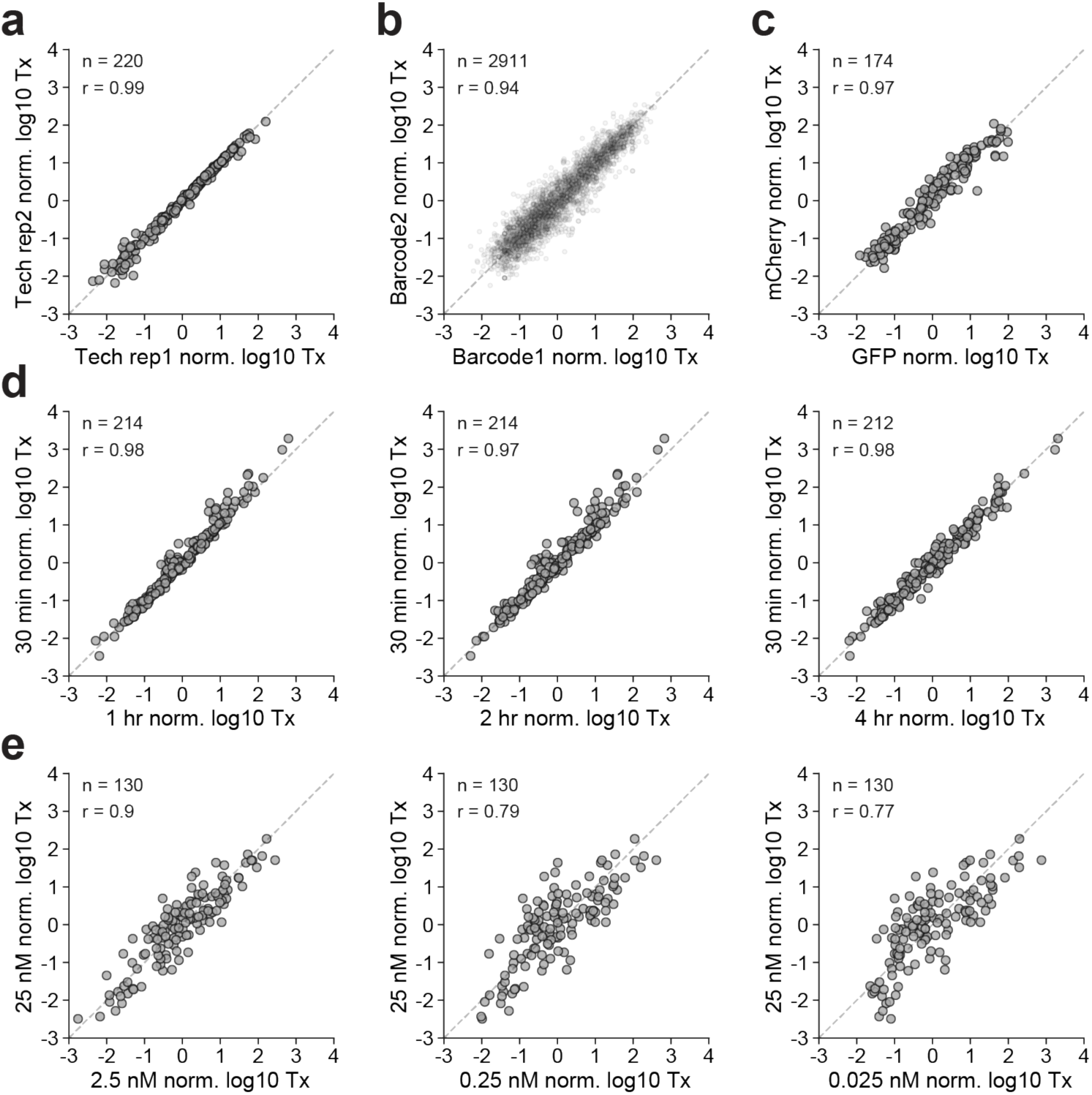
Robustness of DRAFTS transcriptional measurements in *E. coli* cell-free expression system. Comparison of transcriptional profiles (Tx) of regulatory sequences across **(a)** technical replicates, **(b)** alternate barcodes, **(c)** alternate reporter genes, **(d)** reaction times, and **(e)** input DNA concentrations. All measurements except **(a)** are based on merged counts from two technical replicates. Sample sizes (n) and Pearson correlation coefficients (r) can be found in each plot.

**Supplementary Figure S3.**
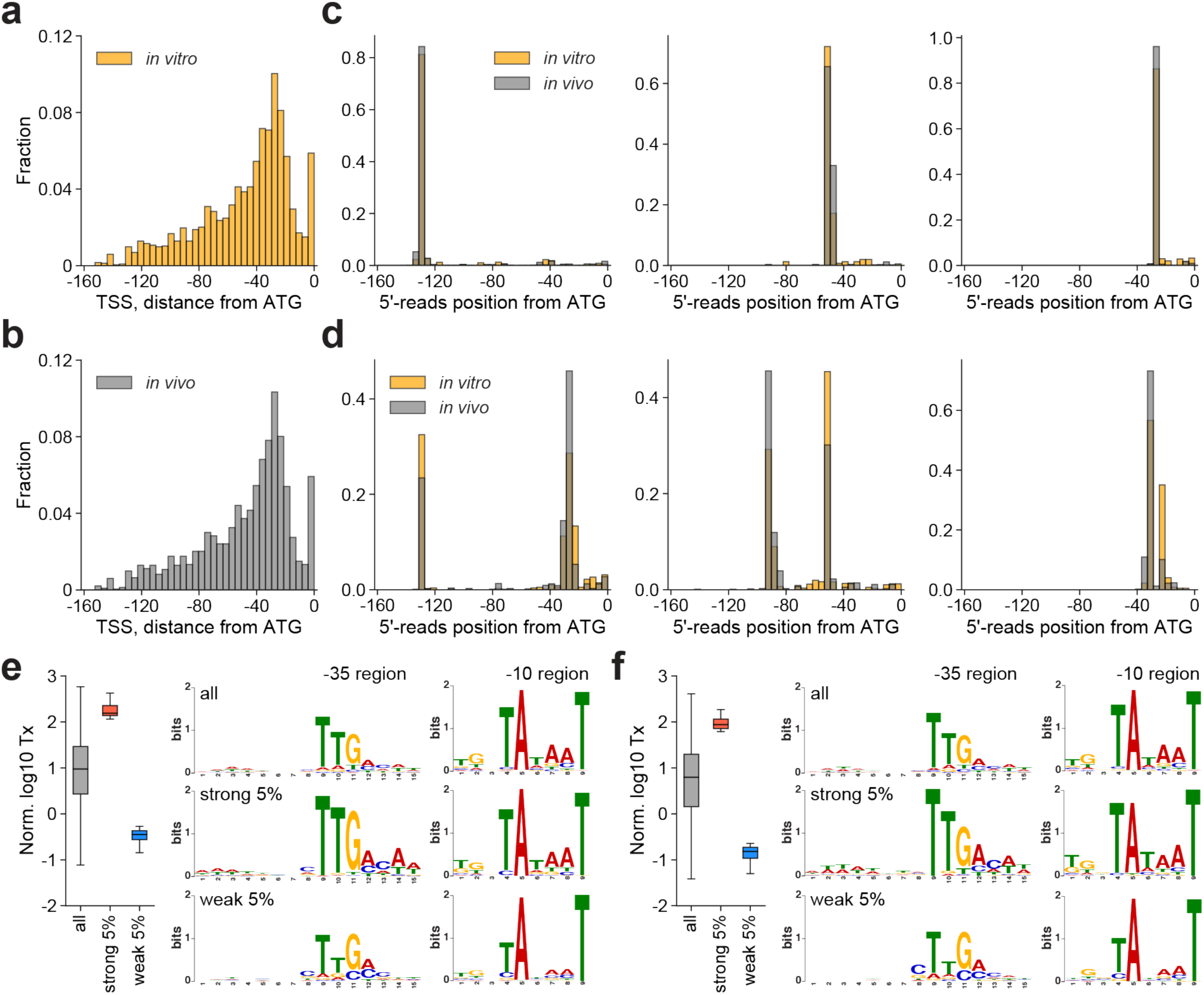
Transcription start site and motif analysis of library regulatory sequences. Primary TSS distributions **(a)** *in vitro* and **(b)** *in vivo*, with example read position distributions for example regulatory sequences containing **(c)** a single and **(d)** two TSSs. Motif analysis on all, strong 5%, and weak 5% promoters from **(e)** *in vitro* or **(f)** *in vivo* measurements gives similar sigma70 motifs enriched for each promoter group.

**Supplementary Figure S4.**
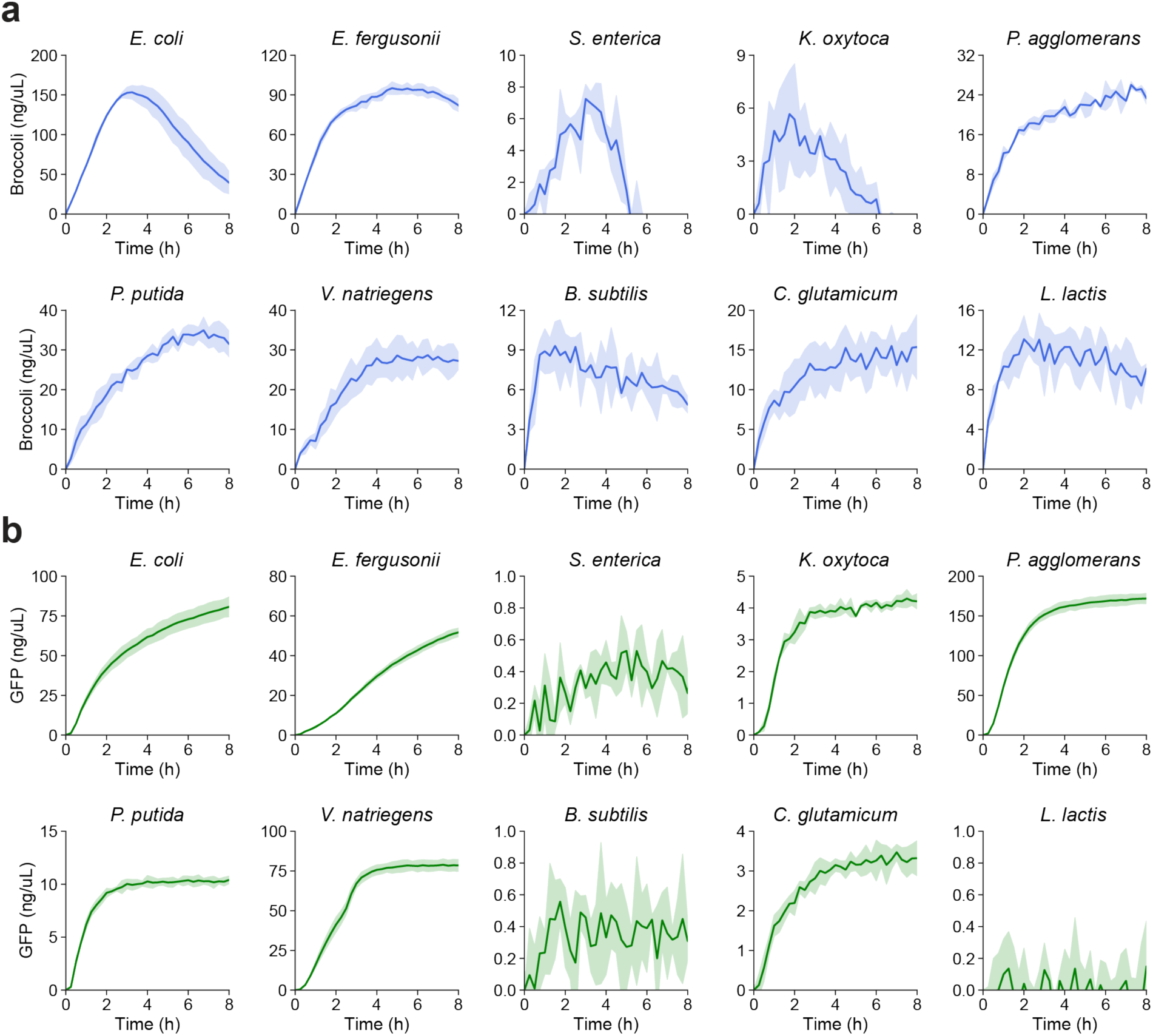
Transcriptional and translational yields from diverse cell-free expression systems. **(a)** Broccoli and **(b)** GFP expression over 8 hours. For Broccoli expression, 12.5 nM of TOPO-F30-Broccoli plasmid was used as a template (50 nM for *L. lactis*). For GFP expression, 25 nM of p70a-deGFP plasmid was used as a template. Shaded regions represent standard deviation of three biological replicates.

**Supplementary Figure S5.**
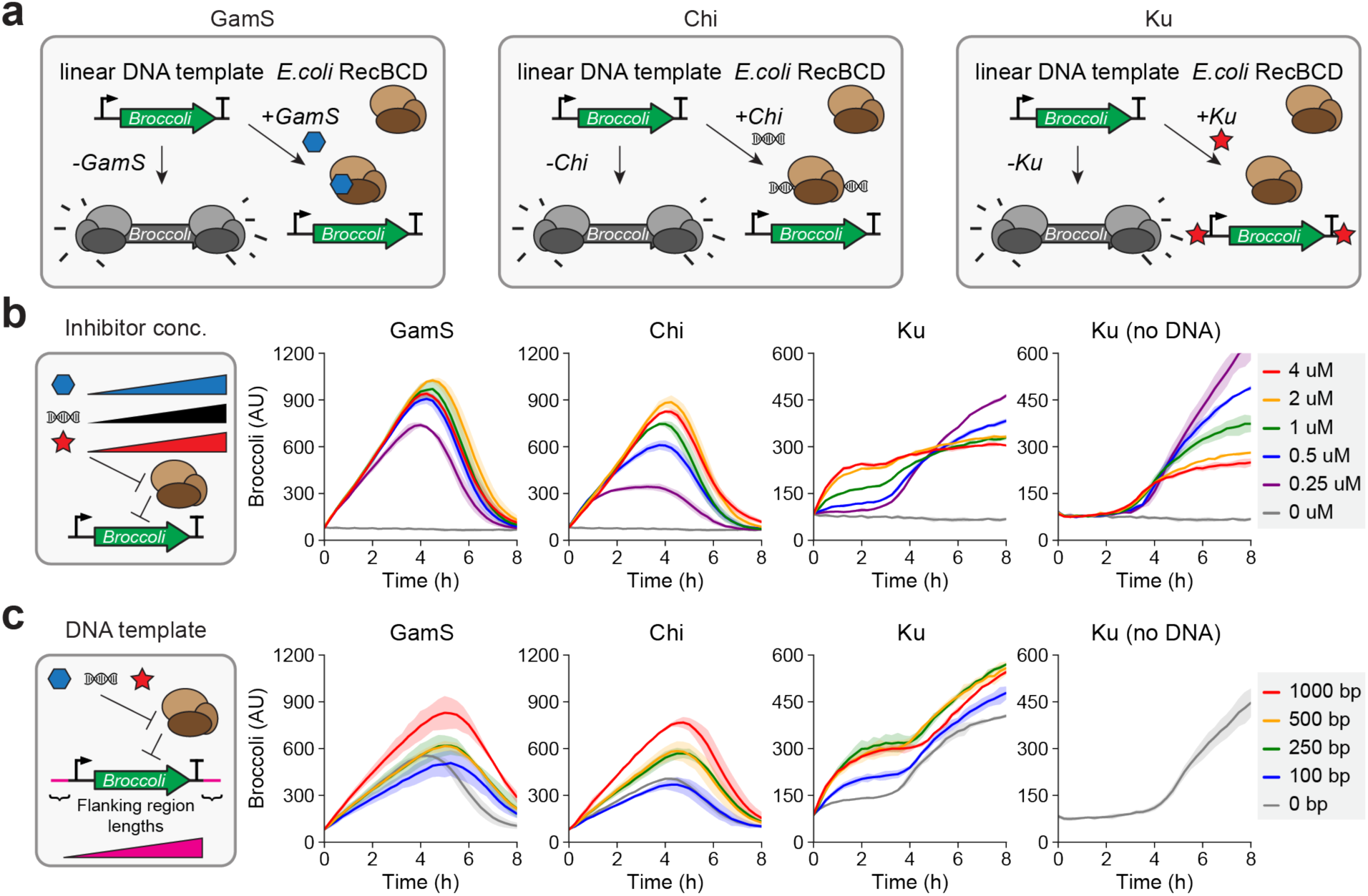
RecBCD inhibitors enhance the stability and activity of linear DNA templates in *E. coli* cell-free expression systems. **(a)** Nuclease inhibiting mechanisms of GamS, Chi, and Ku. Unlike GamS and Chi, Ku from non-homologous end joining (NHEJ) pathway of *Mycobacterium tuberculosis* binds to ends of the linear DNA template, blocking access of nuclease complex to the DNA template. **(b)** Various concentrations of RecBCD inhibitors were tested. **(c)** Examination of flanking sequence length influence on linear DNA stability in presence of three RecBCD inhibitors. Negative control experiments were performed for Ku without DNA template to show fluorescence signals after 3-4 hours of cell-free reaction with Ku is non-specific signals. Commercial *E. coli* cell-free expression system and GamS protein (Arbor Bioscience) were used for all plots shown. Shaded regions represent standard deviation of two biological replicates.

**Supplementary Figure S6.**
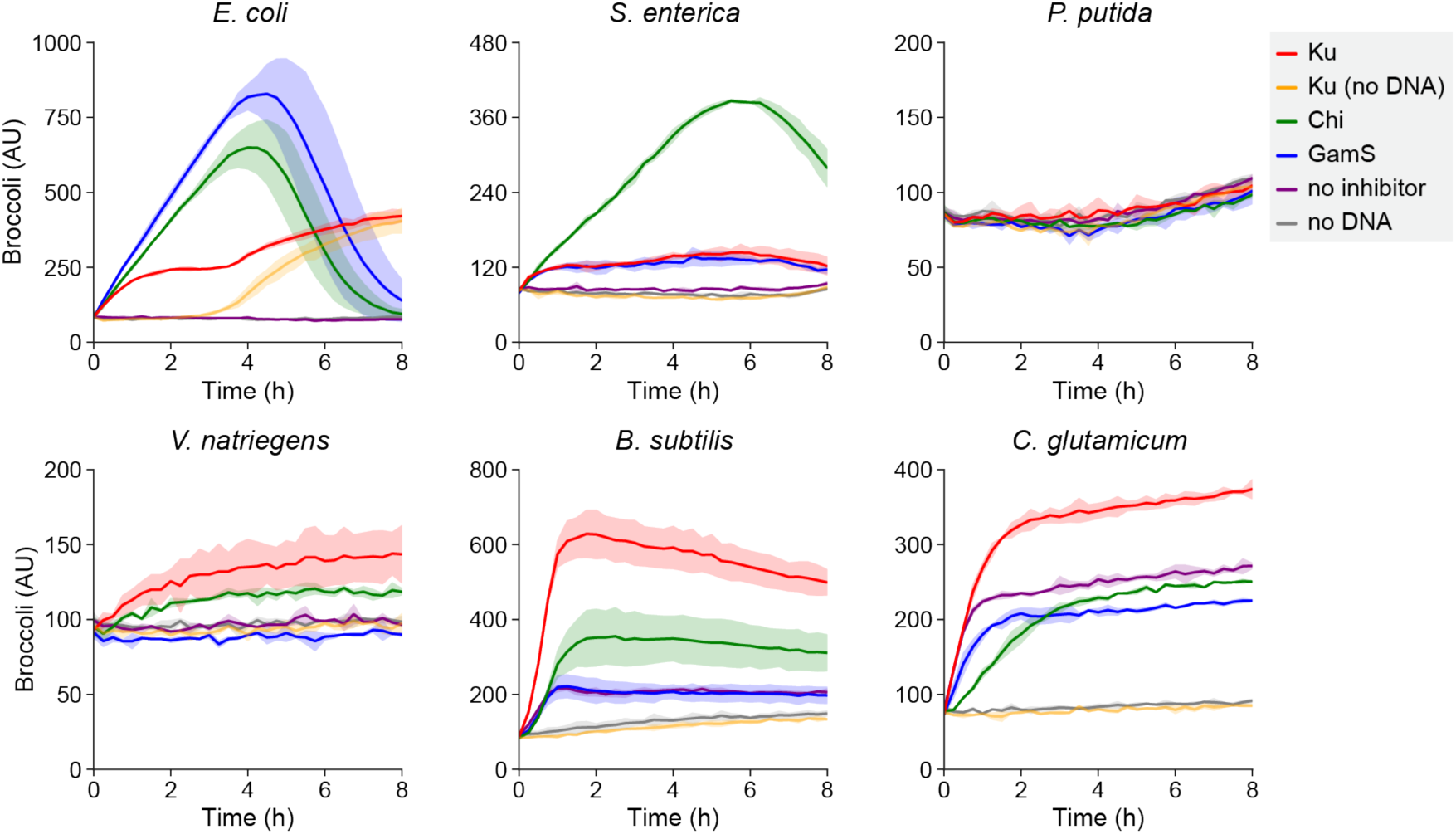
Transcription of Broccoli from a linear DNA template in diverse cell-free expression systems. While GamS and Chi sequence show strong specificity against RecBCD complex from *E. coli* or from closely related bacterial species, Ku could protect the linear DNA template in a broader range of bacterial species, except in *P. putida*, where none of the nuclease inhibitors worked for protection. Commercial *E. coli* cell-free expression system and GamS protein (Arbor Bioscience) were used for *E. coli* plot shown. Shaded regions represent standard deviation of two biological replicates.

**Supplementary Figure S7.**
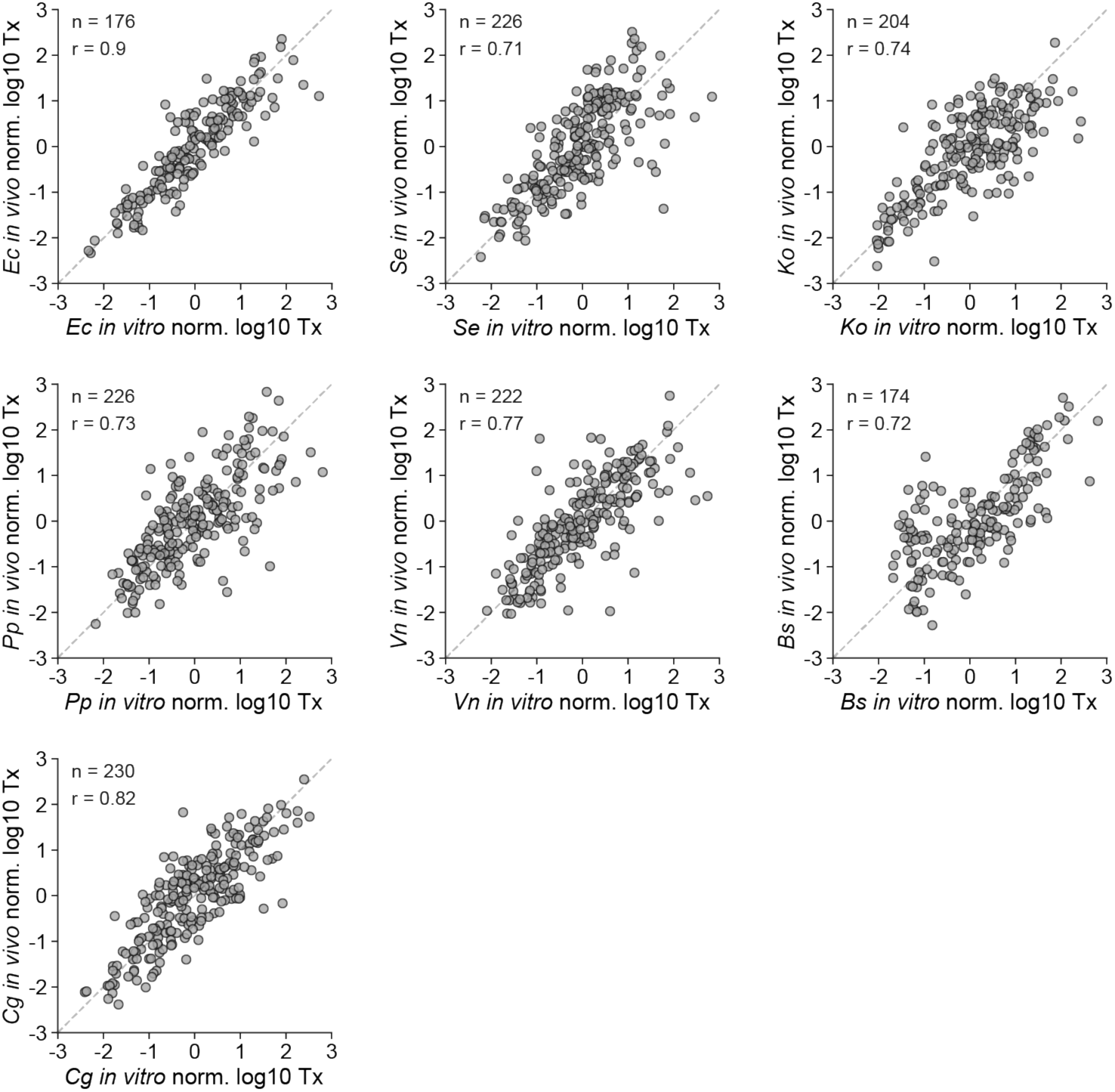
Comparison of *in vitro* and *in vivo* transcriptional measurements in 7 bacterial species. Transcriptional profile (Tx) correlations between *in vivo* and *in vitro* measurements in *E. coli, S. enterica, K. oxytoca, P. putida, V. natriegens, B. subtilis*, and *C. glutamicum*. All transcriptional measurements were made using template-switching adaptor ligation. Sample sizes (n) and Pearson correlation coefficients (r) can be found in each plot.

**Supplementary Figure S8.**
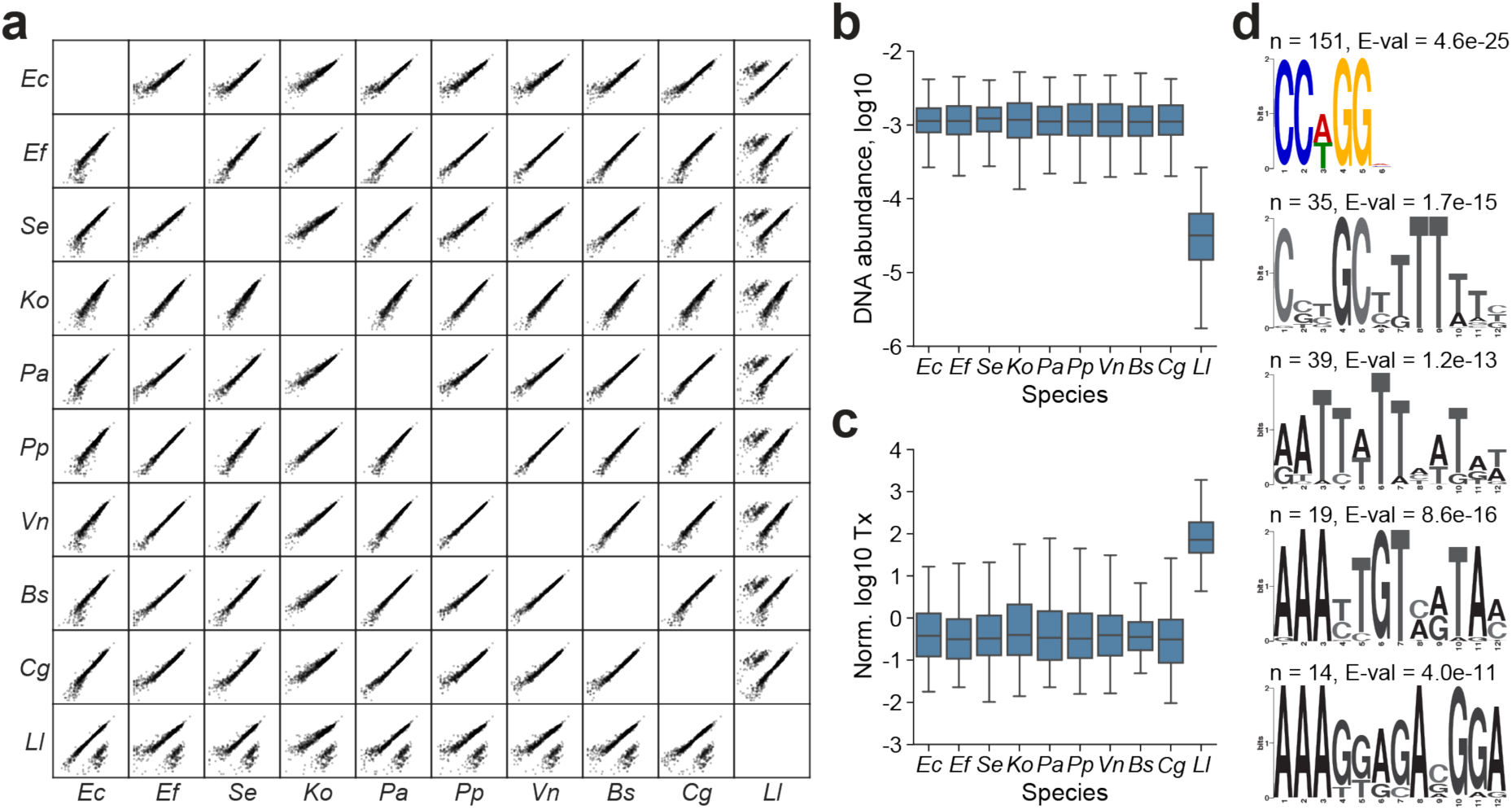
Examination of library DNA abundances identifies sequences subject to restriction enzyme digest in *Lactococcus lactis*. **(a)** Correlation of construct abundances from DNA amplicon sequences after 30-minute incubation in cell extracts from 10 species. **(b)** Box plots of DNA abundance distributions and **(c)** artificially inflated transcriptional activity (Tx) calculations for 182 regulatory sequences that are depleted by >2-fold in *L. lactis*. **(d)** Enriched motifs in *L. lactis* depleted sequences with the number of sequences (n) and motif E-values from MEME listed above each logo. The CCNGG motif had the most hits (151 sites from 182 sequences, 83%) and corresponded to a restriction enzyme (ScrFIR) found in *L. lactis* genome. All others (grayed out) had no putative matching enzyme and had low hit counts.

**Supplementary Figure S9.**
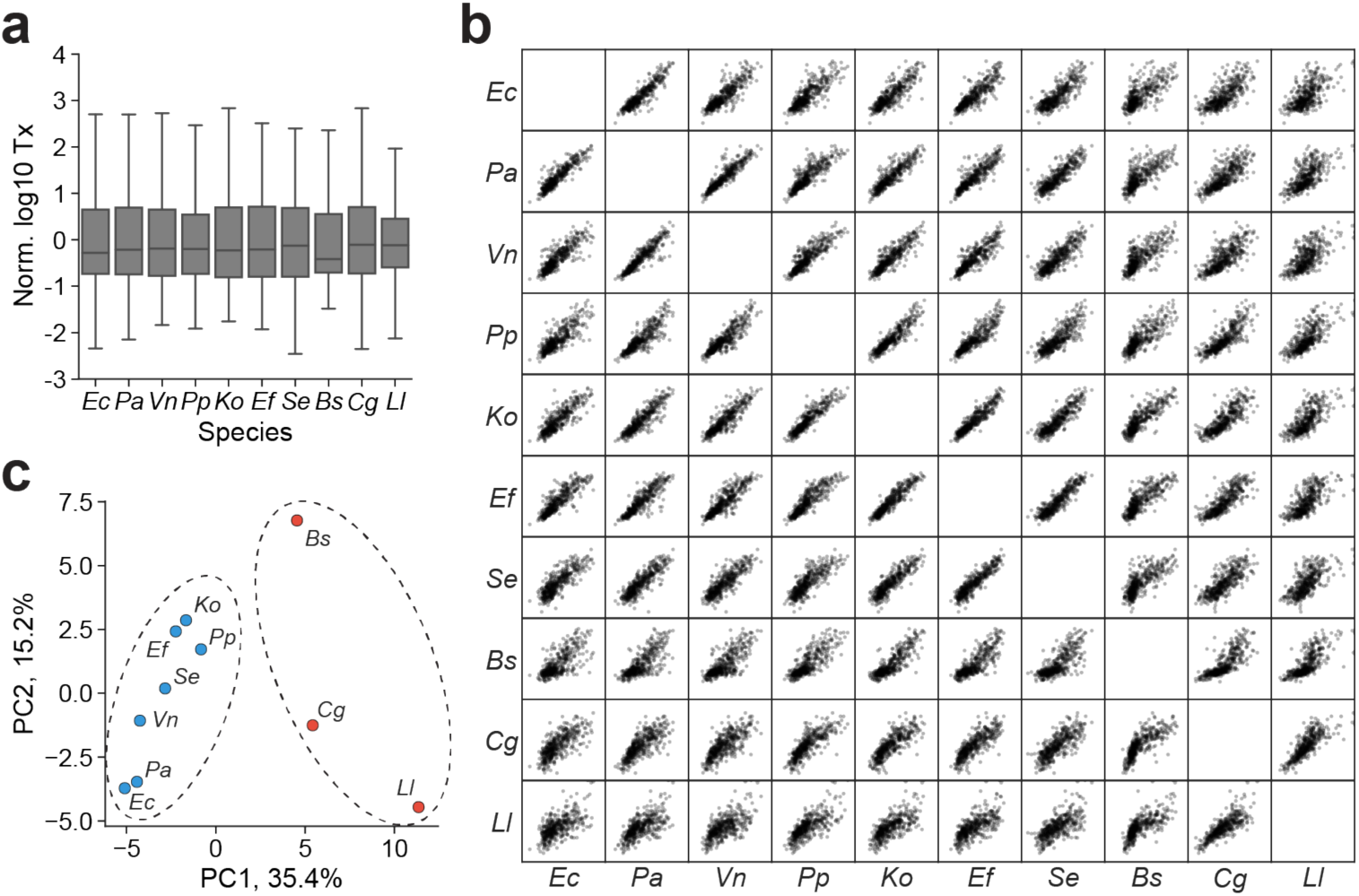
Comparisons of RS1383 transcriptional profiles across 10 bacterial species using DRAFTS. **(a)** Box plot of transcriptional activity (Tx) distributions for each species. **(b)** Scatter plot matrix of *in vitro* transcriptional activity comparisons for 10 species. **(c)** Principal components analysis of transcriptional profile similarity for 10 species.

**Supplementary Figure S10.**
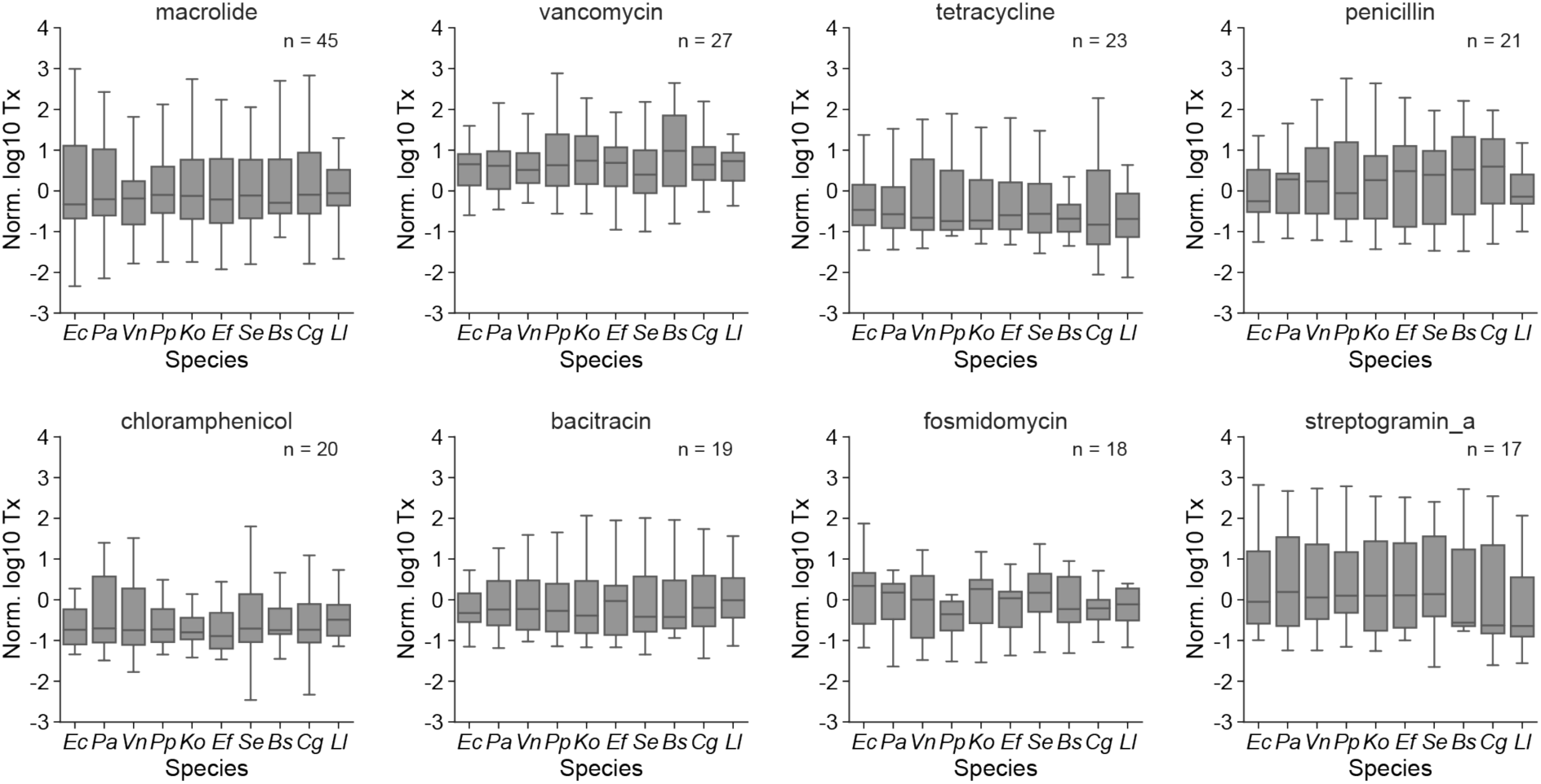
Functional category of regulatory sequence gene origin does not influence activity. Box plots of transcriptional activity (Tx) in 10 species for RS1383 antibiotic resistance gene regulatory sequences, grouped by resistance gene class.

**Supplementary Figure S11.**
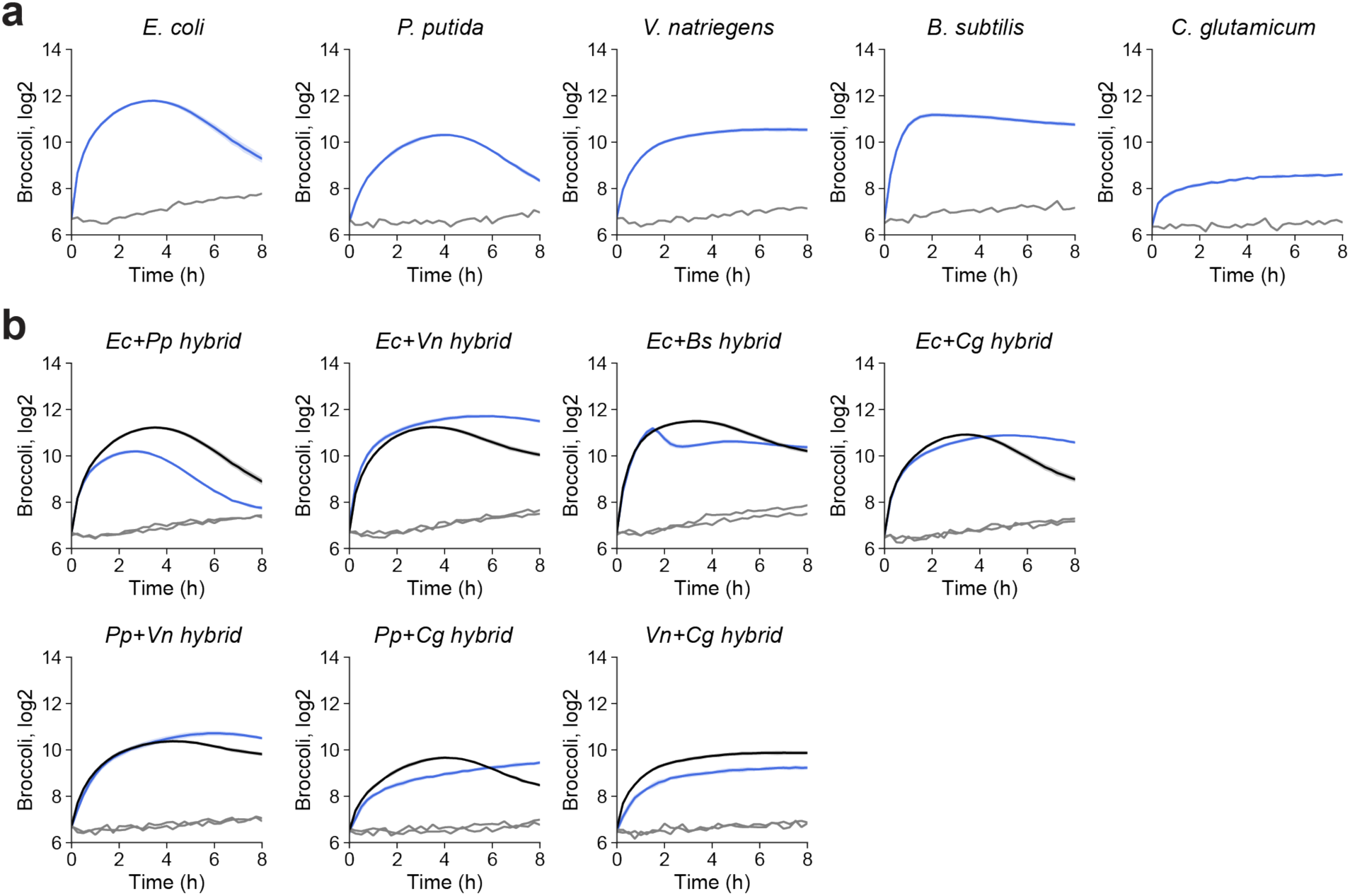
Transcriptional activities from dual-species hybrid lysates. **(a)** Time course profile of Broccoli transcription in 5 single-species source lysates. **(b)** Time course profile of Broccoli transcription in 7 dual-species hybrid lysates. Blue: Observed Broccoli intensity, Grey: Negative control without DNA template, Black: Broccoli intensity predicted by an additive model summing Broccoli intensities from constituent single-species lysates. Shaded regions represent standard deviation of three biological replicates.

**Supplementary Figure S12.**
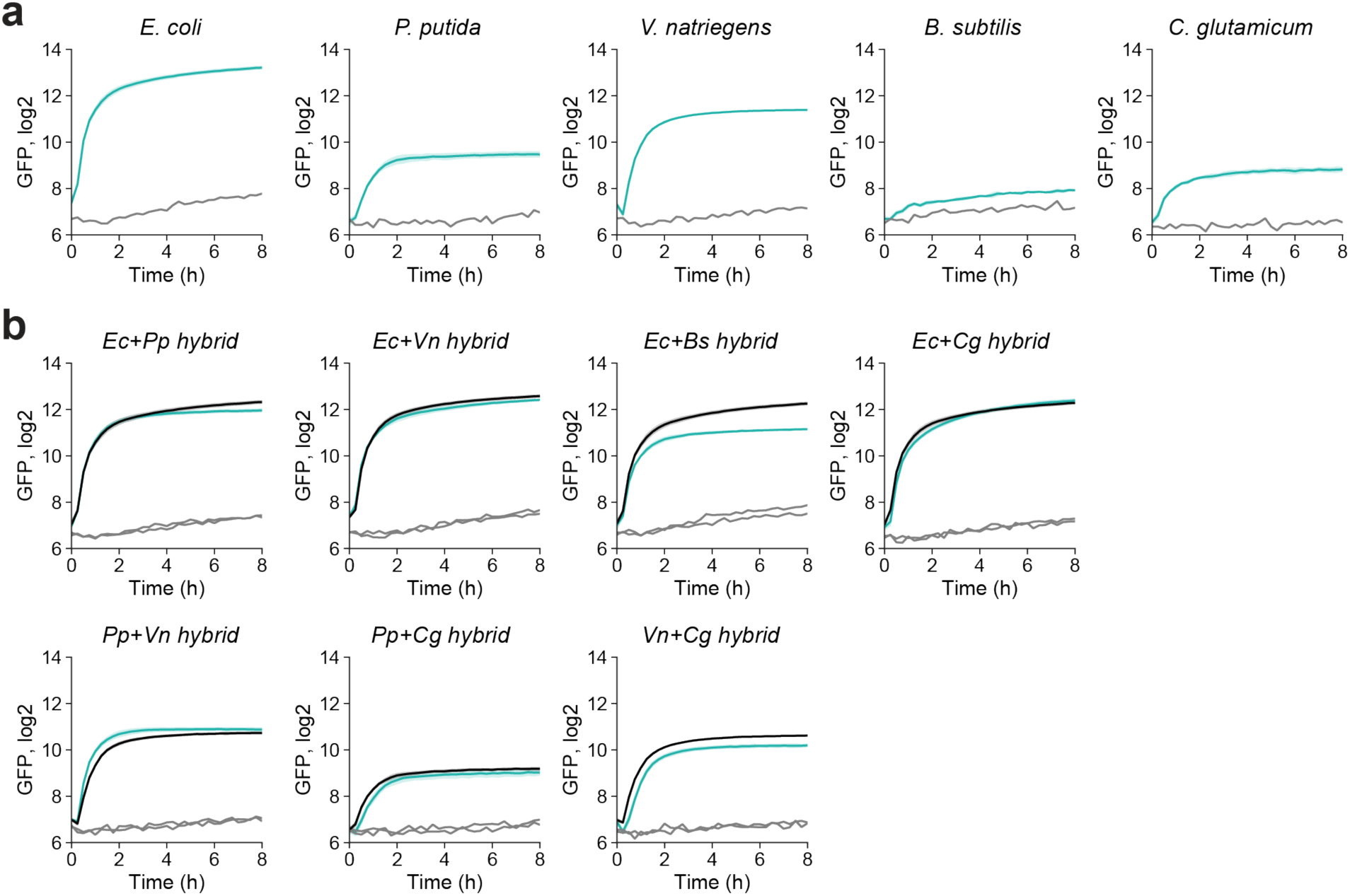
Translational activities from dual-species hybrid lysates. **(a)** Time course profile of GFP translation in 5 single-species source lysates. **(b)** Time course profile of GFP translation in 7 dual-species hybrid lysates. Green: Observed GFP intensity, Grey: Negative control without DNA template, Black: GFP intensity predicted by an additive model summing GFP intensities from constituent single-species lysates. Shaded regions represent standard deviation of three biological replicates.

## References

1 Swartz, J. R. Transforming Biochemical Engineering with Cell-Free Biology. Aiche J 58, 5–13, doi:10.1002/aic.13701 (2012).

2 Shimizu, Y. et al. Cell-free translation reconstituted with purified components. Nat Biotechnol 19, 751–755, doi:10.1038/90802 (2001).

3 Wang, H. H. et al. Multiplexed in vivo His-tagging of enzyme pathways for in vitro single-pot multienzyme catalysis. ACS Synth Biol 1, 43–52, doi:10.1021/sb3000029 (2012).

4 Villarreal, F. et al. Synthetic microbial consortia enable rapid assembly of pure translation machinery. Nat Chem Biol 14, 29–35, doi:10.1038/nchembio.2514 (2018).

5 Garamella, J., Marshall, R., Rustad, M. & Noireaux, V. The All E. coli TX-TL Toolbox 2.0: A Platform for Cell-Free Synthetic Biology. ACS Synth Biol 5, 344–355, doi:10.1021/acssynbio.5b00296 (2016).

6 Jewett, M. C., Fritz, B. R., Timmerman, L. E. & Church, G. M. In vitro integration of ribosomal RNA synthesis, ribosome assembly, and translation. Mol Syst Biol 9, 678, doi:10.1038/msb.2013.31 (2013).

7 Calhoun, K. A., & Swartz, J. R. Energizing cell-free protein synthesis with glucose metabolism. Biotechnol Bioeng 90, 606–613, doi:10.1002/bit.20449 (2005).

8 Jewett, M. C., Calhoun, K. A., Voloshin, A., Wuu, J. J. & Swartz, J. R. An integrated cell-free metabolic platform for protein production and synthetic biology. Mol Syst Biol 4, 220, doi:10.1038/msb.2008.57 (2008).

9 Calhoun, K. A. & Swartz, J. R. Energy systems for ATP regeneration in cell-free protein synthesis reactions. Methods Mol Biol 375, 3–17, doi:10.1007/978-1-59745-388-2_1 (2007).

10 Maini, R., Umemoto, S. & Suga, H. Ribosome-mediated synthesis of natural product-like peptides via cell-free translation. Curr Opin Chem Biol 34, 44–52, doi:10.1016/j.cbpa.2016.06.006 (2016).

11 Dudley, Q. M., Karim, A. S. & Jewett, M. C. Cell-free metabolic engineering: biomanufacturing beyond the cell. Biotechnol J 10, 69–82, doi:10.1002/biot.201400330 (2015).

12 Martin, R. W. et al. Cell-free protein synthesis from genomically recoded bacteria enables multisite incorporation of noncanonical amino acids. Nat Commun 9, 1203, doi:10.1038/s41467-018-03469-5 (2018).

13 Jaroentomeechai, T. et al. Single-pot glycoprotein biosynthesis using a cell-free transcription-translation system enriched with glycosylation machinery. Nat Commun 9, 2686, doi:10.1038/s41467-018-05110-x (2018).

14 Shin, J., Jardine, P. & Noireaux, V. Genome replication, synthesis, and assembly of the bacteriophage T7 in a single cell-free reaction. ACS Synth Biol 1, 408–413, doi:10.1021/sb300049p (2012).

15 Rustad M, E. A., Jardine P, Noireaux V. Cell-free TXTL synthesis of infectious bacteriophage T4 in a single test tube reaction. Synthetic Biology 3, doi:10.1093/synbio/ysy002 (2018).

16 Pardee, K. et al. Paper-based synthetic gene networks. Cell 159, 940–954, doi:10.1016/j.cell.2014.10.004 (2014).

17 Takahashi, M. K. et al. A low-cost paper-based synthetic biology platform for analyzing gut microbiota and host biomarkers. Nat Commun 9, 3347, doi:10.1038/s41467-018-05864-4 (2018).

18 Kwon, Y. C. & Jewett, M. C. High-throughput preparation methods of crude extract for robust cell-free protein synthesis. Sci Rep-Uk 5, doi:ARTN 8663 10.1038/srep08663 (2015).

19 Sun, Z. Z. et al. Protocols for Implementing an Escherichia coli Based TX-TL Cell-Free Expression System for Synthetic Biology. Jove-J Vis Exp, doi:UNSP e50762 10.3791/50762 (2013).

20 Takahashi, M. K. et al. Rapidly characterizing the fast dynamics of RNA genetic circuitry with cell-free transcription-translation (TX-TL) systems. ACS Synth Biol 4, 503–515, doi:10.1021/sb400206c (2015).

21 Hodgman, C. E. & Jewett, M. C. Cell-free synthetic biology: thinking outside the cell. Metab Eng 14, 261–269, doi:10.1016/j.ymben.2011.09.002 (2012).

22 Marshall, R. et al. Rapid and Scalable Characterization of CRISPR Technologies Using an E. coli Cell-Free Transcription-Translation System. Mol Cell 69, 146–157 e143, doi:10.1016/j.molcel.2017.12.007 (2018).

23 Borkowski, O. et al. Cell-free prediction of protein expression costs for growing cells. Nat Commun 9, 1457, doi:10.1038/s41467-018-03970-x (2018).

24 Siegal-Gaskins, D., Tuza, Z. A., Kim, J., Noireaux, V. & Murray, R. M. Gene circuit performance characterization and resource usage in a cell-free “breadboard”. ACS Synth Biol 3, 416–425, doi:10.1021/sb400203p (2014).

25 Chappell, J., Jensen, K. & Freemont, P. S. Validation of an entirely in vitro approach for rapid prototyping of DNA regulatory elements for synthetic biology. Nucleic Acids Res 41, 3471–3481, doi:10.1093/nar/gkt052 (2013).

26 Goodman, D. B., Church, G. M. & Kosuri, S. Causes and effects of N-terminal codon bias in bacterial genes. Science 342, 475–479, doi:10.1126/science.1241934 (2013).

27 Kosuri, S. et al. Composability of regulatory sequences controlling transcription and translation in Escherichia coli. Proc Natl Acad Sci U S A 110, 14024–14029, doi:10.1073/pnas.1301301110 (2013).

28 Kinney, J. B., Murugan, A., Callan, C. G., Jr. & Cox, E. C. Using deep sequencing to characterize the biophysical mechanism of a transcriptional regulatory sequence. Proc Natl Acad Sci U S A 107, 9158–9163, doi:10.1073/pnas.1004290107 (2010).

29 Patwardhan, R. P. et al. High-resolution analysis of DNA regulatory elements by synthetic saturation mutagenesis. Nat Biotechnol 27, 1173–1175, doi:10.1038/nbt.1589 (2009).

30 Vvedenskaya, I. O. et al. Massively Systematic Transcript End Readout, “MASTER”: Transcription Start Site Selection, Transcriptional Slippage, and Transcript Yields. Mol Cell 60, 953–965, doi:10.1016/j.molcel.2015.10.029 (2015).

31 Liu, H. & Deutschbauer, A. M. Rapidly moving new bacteria to model-organism status. Curr Opin Biotechnol 51, 116–122, doi:10.1016/j.copbio.2017.12.006 (2018).

32 Des Soye, B. J., Davidson, S. R., Weinstock, M. T., Gibson, D. G. & Jewett, M. C. Establishing a High-Yielding Cell-Free Protein Synthesis Platform Derived from Vibrio natriegens. ACS Synth Biol, doi:10.1021/acssynbio.8b00252 (2018).

33 Li, J., Wang, H., Kwon, Y. C. & Jewett, M. C. Establishing a high yielding streptomyces-based cell-free protein synthesis system. Biotechnol Bioeng 114, 1343–1353, doi:10.1002/bit.26253 (2017).

34 Kelwick, R., Webb, A. J., MacDonald, J. T. & Freemont, P. S. Development of a Bacillus subtilis cell-free transcription-translation system for prototyping regulatory elements. Metab Eng 38, 370–381, doi:10.1016/j.ymben.2016.09.008 (2016).

35 Moore, S. J. et al. Rapid acquisition and model-based analysis of cell-free transcription-translation reactions from nonmodel bacteria. Proc Natl Acad Sci U S A, doi:10.1073/pnas.1715806115 (2018).

36 Moore, S. J., Lai, H. E., Needham, H., Polizzi, K. M. & Freemont, P. S. Streptomyces venezuelae TX-TL - a next generation cell-free synthetic biology tool. Biotechnol J 12, doi:10.1002/biot.201600678 (2017).

37 Johns, N. I. et al. Metagenomic mining of regulatory elements enables programmable species-selective gene expression. Nature Methods 15, 323-+, doi:10.1038/nmeth.4633 (2018).

38 Swank, Z. L. N; Maerkl, SJ. Cell-free gene regulatory network engineering with synthetic transcription factors. *BioRxiv*, doi:https://doi.org/10.1101/407999 (2018).

39 Silverman, A. K.-L. N; Lucks, JB; Jewett, MC. Deconstructing cell-free extract preparation for in vitro activation of transcriptional genetic circuitry. *BioRxiv*, doi:https://doi.org/10.1101/411785 (2018).

40 Filonov, G. S., Moon, J. D., Svensen, N. & Jaffrey, S. R. Broccoli: rapid selection of an RNA mimic of green fluorescent protein by fluorescence-based selection and directed evolution. J Am Chem Soc 136, 16299–16308, doi:10.1021/ja508478x (2014).

41 Marshall, R., Maxwell, C. S., Collins, S. P., Beisel, C. L. & Noireaux, V. Short DNA containing chi sites enhances DNA stability and gene expression in E. coli cell-free transcription-translation systems. Biotechnol Bioeng 114, 2137–2141, doi:10.1002/bit.26333 (2017).

42 Sinha, K. M., Unciuleac, M. C., Glickman, M. S. & Shuman, S. AdnAB: a new DSB-resecting motor-nuclease from mycobacteria. Genes Dev 23, 1423–1437, doi:10.1101/gad.1805709 (2009).

43 Yeung, E. et al. Biophysical Constraints Arising from Compositional Context in Synthetic Gene Networks. Cell Syst 5, 11–24 e12, doi:10.1016/j.cels.2017.06.001 (2017).

44 Twomey, D. P., Gabillet, N., Daly, C. & Fitzgerald, G. F. Molecular characterization of the restriction endonuclease gene (scrFIR) associated with the ScrFI restriction/modification system from Lactococcus lactis subsp. cremoris UC503. Microbiology 143 ( Pt 7), 2277–2286, doi:10.1099/00221287-143-7-2277 (1997).

45 Urtecho, G., Tripp, A. D., Insigne, K., Kim, H. & Kosuri, S. Systematic Dissection of Sequence Elements Controlling sigma70 Promoters Using a Genomically-Encoded Multiplexed Reporter Assay in E. coli. Biochemistry, doi:10.1021/acs.biochem.7b01069 (2018).

46 Rhodius, V. A. & Mutalik, V. K. Predicting strength and function for promoters of the Escherichia coli alternative sigma factor, sigmaE. Proc Natl Acad Sci U S A 107, 2854–2859, doi:10.1073/pnas.0915066107 (2010).

47 Belliveau, N. M. et al. Systematic approach for dissecting the molecular mechanisms of transcriptional regulation in bacteria. Proc Natl Acad Sci U S A 115, E4796-E4805, doi:10.1073/pnas.1722055115 (2018).

48 Johns, N. I., Blazejewski, T., Gomes, A. L. & Wang, H. H. Principles for designing synthetic microbial communities. Curr Opin Microbiol 31, 146–153, doi:10.1016/j.mib.2016.03.010 (2016).

49 Iyer, L. M., Koonin, E. V. & Aravind, L. Evolution of bacterial RNA polymerase: implications for large-scale bacterial phylogeny, domain accretion, and horizontal gene transfer. Gene 335, 73–88, doi:10.1016/j.gene.2004.03.017 (2004).

50 Noireaux, V., Maeda, Y. T. & Libchaber, A. Development of an artificial cell, from self-organization to computation and self-reproduction. Proc Natl Acad Sci U S A 108, 3473–3480, doi:10.1073/pnas.1017075108 (2011).

51 Chen, Y. et al. Tuning the dynamic range of bacterial promoters regulated by ligand-inducible transcription factors. Nat Commun 9, 64, doi:10.1038/s41467-017-02473-5 (2018).

52 Sharon, E. et al. Inferring gene regulatory logic from high-throughput measurements of thousands of systematically designed promoters. Nat Biotechnol 30, 521–530, doi:10.1038/nbt.2205 (2012).

53 Rhodius, V. A. et al. Design of orthogonal genetic switches based on a crosstalk map of sigmas, anti-sigmas, and promoters. Mol Syst Biol 9, 702, doi:10.1038/msb.2013.58 (2013).

54 Stanton, B. C. et al. Genomic mining of prokaryotic repressors for orthogonal logic gates. Nat Chem Biol 10, 99–105, doi:10.1038/nchembio.1411 (2014).

55 Zong, D. M. et al. Predicting Transcriptional Output of Synthetic Multi-input Promoters. ACS Synth Biol 7, 1834–1843, doi:10.1021/acssynbio.8b00165 (2018).

56 Gorochowski, T. E. et al. Genetic circuit characterization and debugging using RNA-seq. Mol Syst Biol 13, 952, doi:10.15252/msb.20167461 (2017).

57 Ingolia, N. T., Ghaemmaghami, S., Newman, J. R. & Weissman, J. S. Genome-wide analysis in vivo of translation with nucleotide resolution using ribosome profiling. Science 324, 218–223, doi:10.1126/science.1168978 (2009).

58 Li, G. W., Burkhardt, D., Gross, C. & Weissman, J. S. Quantifying absolute protein synthesis rates reveals principles underlying allocation of cellular resources. Cell 157, 624–635, doi:10.1016/j.cell.2014.02.033 (2014).

59 Hodgman, C. E. & Jewett, M. C. Optimized extract preparation methods and reaction conditions for improved yeast cell-free protein synthesis. Biotechnol Bioeng 110, 2643–2654, doi:10.1002/bit.24942 (2013).

60 Sawasaki, T., Morishita, R., Gouda, M. D. & Endo, Y. Methods for high-throughput materialization of genetic information based on wheat germ cell-free expression system. Methods Mol Biol 375, 95–106, doi:10.1007/978-1-59745-388-2_5 (2007).

61 Ezure, T. et al. Cell-free protein synthesis system prepared from insect cells by freeze-thawing. Biotechnol Prog 22, 1570–1577, doi:10.1021/bp060110v (2006).

62 Mikami, S., Masutani, M., Sonenberg, N., Yokoyama, S. & Imataka, H. An efficient mammalian cell-free translation system supplemented with translation factors. Protein Expres Purif 46, 348–357, doi:10.1016/j.pep.2005.09.021 (2006).

63 Martin, R. W. et al. Development of a CHO-Based Cell-Free Platform for Synthesis of Active Monoclonal Antibodies. ACS Synth Biol 6, 1370–1379, doi:10.1021/acssynbio.7b00001 (2017).

64 Liu, B. & Pop, M. ARDB--Antibiotic Resistance Genes Database. Nucleic Acids Res 37, D443–447, doi:10.1093/nar/gkn656 (2009).

65 Chen, L., Xiong, Z., Sun, L., Yang, J. & Jin, Q. VFDB 2012 update: toward the genetic diversity and molecular evolution of bacterial virulence factors. Nucleic Acids Res 40, D641–645, doi:10.1093/nar/gkr989 (2012).

66 Stead, M. B. et al. RNAsnap: a rapid, quantitative and inexpensive, method for isolating total RNA from bacteria. Nucleic Acids Res 40, e156, doi:10.1093/nar/gks680 (2012).

67 Bushnell, B., Rood, J. & Singer, E. BBMerge - Accurate paired shotgun read merging via overlap. PLoS One 12, e0185056, doi:10.1371/journal.pone.0185056 (2017).

68 Zadeh, J. N. et al. NUPACK: Analysis and design of nucleic acid systems. J Comput Chem 32, 170–173, doi:10.1002/jcc.21596 (2011).

69 Bailey, T. L. Discovering novel sequence motifs with MEME. Curr Protoc Bioinformatics Chapter 2, Unit 2 4, doi:10.1002/0471250953.bi0204s00 (2002).

70 Yus, E., Yang, J. S., Sogues, A. & Serrano, L. A reporter system coupled with high-throughput sequencing unveils key bacterial transcription and translation determinants. Nat Commun 8, 368, doi:10.1038/s41467-017-00239-7 (2017).

